# Who takes the risk to forage? Testing consistent inter-individual behavioural variation in wild vervet monkeys (*Chlorocebus pygerythrus*)

**DOI:** 10.64898/2026.06.08.730785

**Authors:** des Pallières Claire Gauquelin, Belli Elena, Aguilera Fanny, Halbwax Michel, Šlipogor Vedrana, van de Waal Erica, Koren Lee, Matas Devorah, Canteloup Charlotte

## Abstract

Risk-taking behaviour permeates daily decision-making across many taxa and has significant impact on fitness outcomes. Previous research finds that individuals display consistent tendencies in their propensity to take risks, often labelled as a “Boldness-Shyness” personality trait. In this experimental study, we investigated repeatability in risk-taking behaviour in 241 wild vervet monkeys by measuring their responses to capture threat (via human-initiated darting procedure) and predation threat (via two predator models). We further examined the influence of socio-demographic, group identity and hormonal factors on risk-taking behaviour. We found that multiple behavioural responses, including the likelihood of approaching a food source under threat, were consistent across contexts. Risk-taking behaviour was influenced by age, sex, dominance rank, and hormonal profiles: juveniles, males, and higher-ranking individuals were more likely to approach food under (perceived) predation risk. Additionally, we observed significant among-group differences, suggesting that individuals within the same social group exhibit similar risk sensitivities, potentially due to social facilitation or shared environmental effects. The findings contribute to our understanding of animal personality and the ecological and social drivers underlying variation in risk-related behaviours.

## Introduction

Risk-taking, broadly defined as the behavioural response to unfamiliar or threatening situations, plays a central role in the lives of animals [1]. Taking or avoiding risks is essential for navigating social hierarchies and dynamics, discovering new territories and ensuring survival during foraging or predator encounters [2]. Perception of risk is therefore likely to have strong effects on many aspects of decision-making such as in evaluating resource quantity, quality, dispersion, distance and visibility during foraging. Foraging decisions also require animals to balance energetic gains against associated risks such as predation [3–6] and within-group competition for food, which can result in injuries or even death [7]. Accurate risk assessment is thus likely to have strong fitness consequences. However, individuals across species—including humans—do not always behave in a fully rational or optimal way when faced with risky decisions [8,9]. Human research has shown that individuals rather markedly differ in their tolerance for risk, and often this appears to reflect certain stable personality traits such as risk-aversion or boldness [10,11]. These consistent inter-individual differences are not unique to humans, with studies in a wide range of taxa from fishes, birds, ungulates, rodents and non-human primates (hereafter primates), showing similar patterns [1,4,12–16].

Boldness is one of the five categories of animal personality traits proposed by Réale and colleagues [17] and is often assessed by observing responses to predator-related behaviours, but also sometimes novel objects or food (see also [18]). Such personality assessments using predator models have been validated in a variety of primate species (e.g., common marmosets (*Callithrix jacchus*): [19,20]; barbary macaques (*Macaca sylvanus*): [21,22]; vervet monkeys (*Chlorocebus pygerythrus*): [23]; chacma baboons (*Papio ursinus*): [24]) with models eliciting ecologically relevant antipredator behaviours and avoidance strategies including elevated vigilance, alarm calls, and avoidance [25]. Predation is one of the major causes of death in primates [26] and individual differences in antipredator strategies may thus contribute greatly to their fitness [6,27]. Interestingly, individual consistency in risk-taking behaviours is higher under high predation risk compared to low predation risk in mouse lemurs (*Microcebus murinus*: [4]), suggesting that individual variation may also be flexible and therefore adaptable to given circumstances or severity of threat.

In social species, risk-taking behaviour does not occur in isolation. Group size, habitat structure, and social composition can all influence vulnerability to predation and shape antipredator strategies [28,29]. Across taxa, individuals in larger or mixed-species groups typically reduce vigilance as they benefit from risk dilution or collective detection [30]. It is therefore essential to understand how social context influences decision-making, especially in highly gregarious taxa such as primates [31]. Despite this, most risk-taking studies have been done in captive and/or solitary settings, relying on small sample sizes (4-24 individuals) and limited demographic characteristics (reviewed in [32]), instead of within natural social group context and ecologically relevant conditions in the wild. Notable exceptions include a recent study by Šlipogor and colleagues [33] who identified consistent inter-individual differences in response to a snake model in wild common marmosets in the Cattinga forests of Brazil. Beyond individual traits, social processes may further shape risk-taking behaviour. Social traditions and group norms may lead individuals within the same group to develop similar behavioural tendencies [34,35]. This pattern has been observed in marmosets, where group membership was shown to predict behavioural similarity in response to a range of experiments exploring Boldness and Exploration, suggesting that social transmission may influence risk preferences alongside individual traits [19,36]. In a recent study, Šlipogor and collaborators [20] compared individual and social testing of common marmosets in captivity and found that whilst the trait ‘Boldness/Exploration’ was consistent across a social or solitary setting, the trait ‘Stress/Activity’ changed depending on the social setting; namely, individuals that expressed lower ‘Stress/Activity’ values when alone showed higher ‘Stress/Activity’ values in the presence of group members, and vice versa; showing that the social versus solitary setting can influence behavioural expression in animals in some contexts. Together, these findings highlight the need for studies that simultaneously examine individual consistency and social influences on risk-taking, particularly in wild populations.

Risk-taking behaviour is also shaped by socio-demographic factors. Age-dependent differences have been reported with younger individuals often exhibiting higher exploration and risk-proneness than adults in chimpanzees (*Pan troglodytes,* [37]) and adult Balinese long-tailed macaque (*Macaca fascicularis fascicularis*) females being more risk-prone than juveniles, while juvenile and older adult males being more risk prone than younger adults [38]. However, no age effect has been found in captive Barbary macaques [21] and other primate species, although this is possibly attributable to small sample sizes in conducted studies [13,39,40]. An age effect has also been reported in wild vervet monkeys in the context of novel object and predator exposure experiments [23]. In this study, individuals were presented with a novel predator model (i.e. a lizard) alongside observations of natural snake encounters to assess consistency in boldness-related behaviours. Behavioural responses such as approaches to the model and inspection of real predators were found to be repeatable across contexts. Notably, subadult males exhibited higher levels of boldness than adult females, including more frequent inspection of natural predators. However, evidence for sex and dominance rank effects on risk-taking in primates remains mixed. A recent study testing captive Tonkean macaques (*Macaca tonkeana*) in a gambling task on touchscreens revealed that middle ranking individuals exhibit reduced risk aversion in the gain but not in the loss domain [41]. However, a general pattern suggests that males and higher-ranking individuals tend to engage more readily in risky or novel situations than females and lower-ranking individuals [1,32,37,42]. These differences are often interpreted through the lens of sex-specific life-history strategies and social roles – for example, greater dispersal tendencies or competitive pressures in males, and higher reproductive costs in females [43,44]. Comparable patterns in humans further suggest that variation in risk-related behaviour may arise from both genetic predispositions and culturally transmitted norms, potentially reflecting deep evolutionary roots [37,45].

Links between risk-taking behaviour and physiological markers have also been reported and are important to consider. In particular, cortisol and testosterone are associated with differences in risk-taking, although these effects are inconsistent across studies, particularly in the extensive human literature [46,47]. Some studies reported a positive correlation between testosterone levels and risk-taking [46], whereas others found a negative correlation [48], no effect [49] or some non-linear effects [50]. Similarly, cortisol, a hormone associated with stress responses, has been linked to higher risk-taking in some contexts, and its effects were more pronounced in men than women ([51,52], but see [53]). Moreover, the “dual hormone hypothesis” posits that testosterone and cortisol interact in such a way that testosterone level is positively related to risk-taking and aggression but only in individuals with low cortisol [54]. Similar research in non-human primates remains limited and results are inconsistent. In one study, captive vervet monkeys with lower baseline cortisol levels exhibited bolder responses to novel objects [55,56] while another study found that common marmosets with higher cortisol levels showed increased exploratory behaviours [57]. Most of the primate studies have been conducted in captive settings such as laboratories and research facilities, likely due to the logistical challenges of collecting physiological samples without disrupting behaviour in the wild [58]. Further research in natural populations is therefore needed to assess whether these relationships hold under ecologically relevant settings.

The present study addresses the gaps highlighted above by investigating individual- and group-level differences in response to risk, alongside associated hormonal profiles, in seven groups of wild vervet monkeys. Vervet monkeys provide an excellent model to address such questions because, on the one hand, they are highly social and cohesive (Cheney & Seyfarth, 1990), display conformity in social and foraging contexts (Kerjean et al., 2024; van de Waal et al., 2013a), and exhibit coordinated group-level antipredator responses [59,60]. On the other hand, they show pronounced inter-individual variation in behavioural tendencies consistent with animal personality traits [23,61–63]. Studying such variation provides valuable insights into traits that influence key ecological and evolutionary processes such as niche expansion, dispersal, and social organisation [17]. We examined risk-taking behaviours across three high-risk foraging scenarios, i.e., feeding under (1) threat of human capture (darting) (2) predation risk from a snake model, (3) predation risk from an eagle model. The darting is a more “uncertain” scenario where the threat is invisible (i.e., the veterinarian and dart gun are hidden within a hide) and irregular (occurs at maximum once to twice per year at different times). Snakes and eagles are natural predators of vervet monkeys and represent distinct aerial and terrestrial predatory threats that elicit different behavioural responses [60]. While snake models are often used in predator-response studies, bird-of-prey models are more rarely used and risk-taking in response to capture-related threats such as darting is particularly underexplored, with only a few studies addressing this type of anthropogenic risk [64,65]. Using these three experimental contexts, we addressed the following questions: i) do individuals display intra-individual and inter-individual behavioural consistency in response to risk?; ii) Do socio-demographic factors (i.e. age, sex and dominance rank) predict risky behaviours?; iii) How do these socio-demographic factors relate to hormonal profiles?; iv) How are hormonal profiles associated with risk-taking behaviours?; and v) Do groups exhibit distinct, group-level behavioural tendencies?

Regarding i), we predicted that individuals would show consistent behavioural tendencies both across contexts and across repeated trials of each context. We further predicted that individuals would be more risk-prone in response to the less predictable darting scenario than to the visible and familiar predation threats of aerial and terrestrial predators. Regarding ii), we expected younger individuals to be less risk-averse than older individuals [4,66], and females to be more risk-averse than males due to differing reproductive strategies and parental investment [37,43], while not making any strong predictions regarding dominance rank. Regarding iii), we predicted that high-ranking individuals would exhibit lower cortisol levels than low-ranking individuals [67], that males would show higher testosterone and lower cortisol than females [55,56,68], and that juveniles would have higher testosterone than adults [55,56]. Regarding iv), we predicted that individuals with higher-than-average testosterone level in hair and lower-than-average cortisol level in hair would exhibit greater risk-taking behaviour [55,56,69]. Finally, for v), we predicted individuals within the same group to show more similar responses than individuals from different groups [34,36]. By integrating behavioural and hormonal measures across multiple ecologically relevant risk contexts and by sampling individuals from seven social groups, this study provides valuable insights into the relative contributions of individual traits, social environment, and context to risk-related decision-making in a wild primate species.

## Results

### Repeatability

Repeatability across trials varied for different behavioural variables and experiment types. The behavioural variable “time spent at the corn” was significantly repeatable according to the LRT test (p ≤ 0.01), with the highest r value for snake (r = 0.42), followed by eagle (r = 0.38), and darting (r = 0.21). The number of “scans within 1 to 5m of the corn” was also significantly repeatable across all experiment types (p ≤ 0.01), and the r value was highest for snake (r = 0.18), then eagle (r = 0.15) and darting (r = 0.09). “Scans in >10m of the corn” was not repeatable either for snake (p = 0.05) or for eagle (p > 0.05), but it was repeatable for darting (p ≤ 0.001). “Alarm call” was repeatable for eagle, with the highest r value (r = 0.46; p ≤ 0.001), and for snake (r = 0.18; p ≤ 0.001). “Approach the model” was repeatable for darting (r = 0.68; p ≤ 0.001), although we could not run bootstraps for this as mentioned earlier (i.e. because of its low occurrence number), and it was not repeatable for snake (r = 0.05; p = 0.05).

Regarding the repeatability across the three experiment types, “time spent at the corn” was repeatable (r = 0.24; p ≤ 0.001), as was “scans within 1 to 5m of the corn” (r = 0.13; ≤ 0.001), and “alarm call” (r = 0.16; p ≤ 0.001).

### Effects of socio-demographic variables and hormonal levels on risky behaviours

#### Comparison of “boldness towards corn” between groups

We found significant differences in “boldness towards the corn” values among different monkey groups (Kruskal-Wallis test: χ^2^ = 36.22, df = 5, p < 0.0001; Fig 1). Dunn’s test for post-hoc pairwise comparisons highlighted that AK showed significantly higher boldness than BD (AK-BD: z = 3.162, p = 0.023) and LT (AK-LT: z = 4.929, p < 0.0001), and that KB also showed higher boldness than BD (BD-KB: z = -3.114, p = 0.028) and LT (KB-LT: z = 4.884, p < 0.0001). We did not find any significant difference in “boldness towards the corn” score between the other groups (AK-CR: z = 2.856; BD-CR: z = 0.028; AK-KB: z = 0.042; CR-KB: z = -2.813; BD-LT: z = 2.381; CR-LT: z = 2.070; AK-NH: z = 2.465; BD-NH: z = - 0.557; CR-NH: z = -0.515; KB-NH: z = 2.420; LT-NH: z = -2.688; p > 0.05).

**Fig 1.**
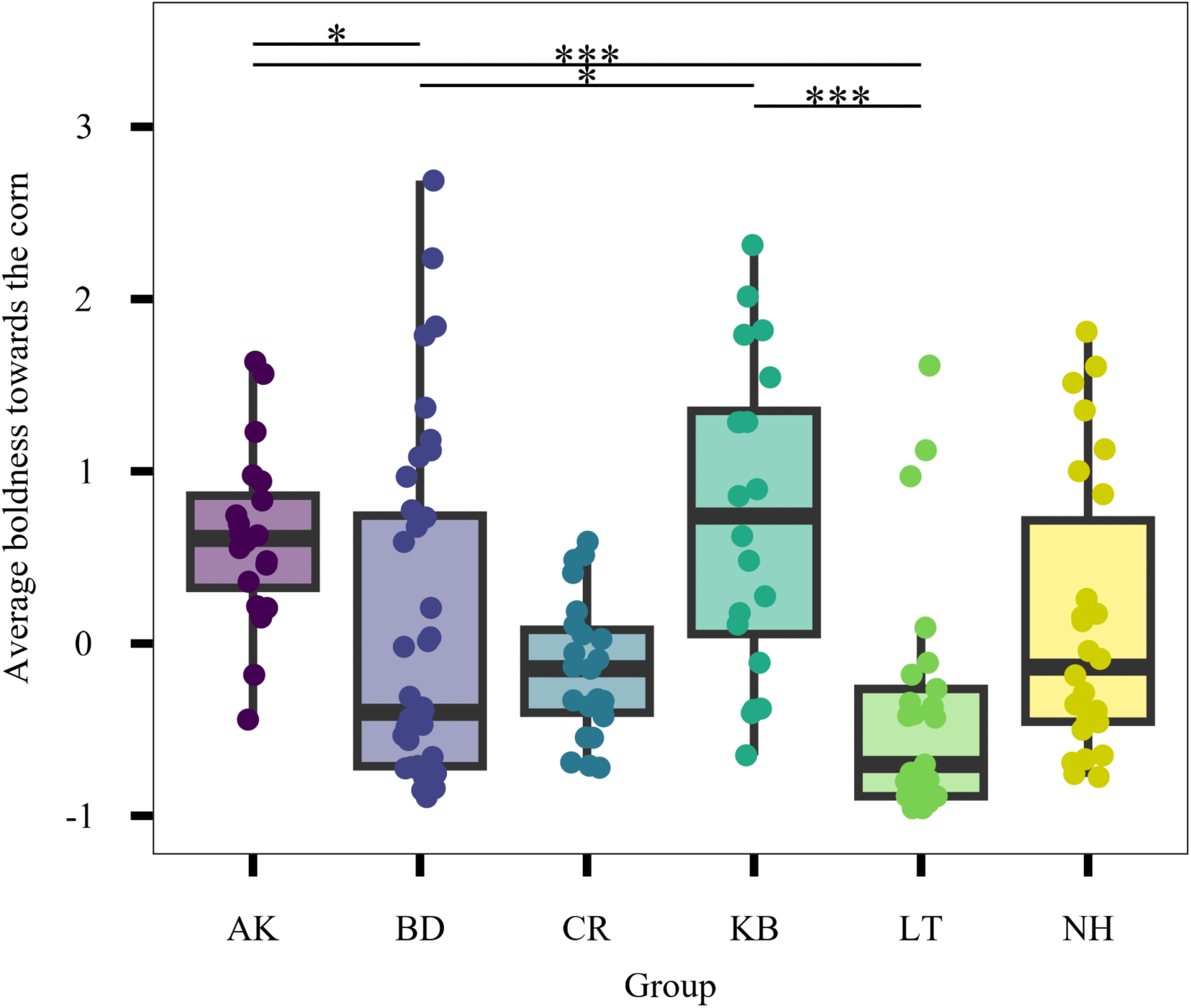
Variation in boldness towards the corn across groups. Boldness towards the corn scores were calculated as the mean PC1 score derived from a principal component analysis combining behavioural responses across darting, snake, and eagle contexts, resulting in one composite score per individual. Higher values indicate greater boldness. Points represent individual monkey boldness towards the corn scores; boxes represent the interquartile range (IQR) with the median shown as the central line; whiskers extend to 1.5 × IQR. * p < 0.05; ** p < 0.01; *** p < 0.001.

#### Effects of age, sex, rank and experiment on locations relative to the corn during scans

In GLMM1, the effect of age was significant (LRT = 5.68; p = 0.017) with juveniles being 1.60 times (60%) more likely to be within 1m of the corn than adults (Estimate = 0.47; SE = 0.20; z = 2.39, p = 0.017; Fig 2a). The effect of sex was significant (LRT = 4.26; p = 0.039) with males being 1.52 times (52%) more likely to stand within 1m of the corn than females (Estimate = 0.42; SE = 0.20; z = 2.09; p = 0.037; Fig 2b). The effect of rank was significant (LRT = 27.21; p <0.0001); with each increase of rank unit associated with 9.7 times higher odds to be at the corn (Estimate = 2.27; SE = 0.43; z = 5.33, p <0.001; Fig 2c). The effect of experiment type was significant (LRT = 19.33; p < 0.0001) with individuals spending less time within 1m of the corn during darting compared to snake (Estimate = -0.22; SE = 0.05; z = -4.36; p < 0.0001; Fig 2d). However, Tukey post hoc contrasts showed no significant difference between darting and eagle (Estimate = -0.16; SE = 0.07; z = -2.23; p > 0.05) and between snake and eagle (Estimate = 0.06; SE = 0.07; z = 0.88; p > 0.05; Fig 2). The random effect of ID nested within group had a high variance of σ² = 1.75 (SD = 1.32) and group alone had a variance of σ² = 0.49 (SD = 0.70). The variance from trial number was small (σ² = 0.04; SD = 0.20). The marginal R² was 0.12, indicating that the fixed effects explained approximately 12% of the variance in “scans within 1m of the corn”. The conditional R² was 0.89, suggesting that the full model—including the random effects of ID nested within group and trial number —explained approximately 89% of the variance.

**Fig 2.**
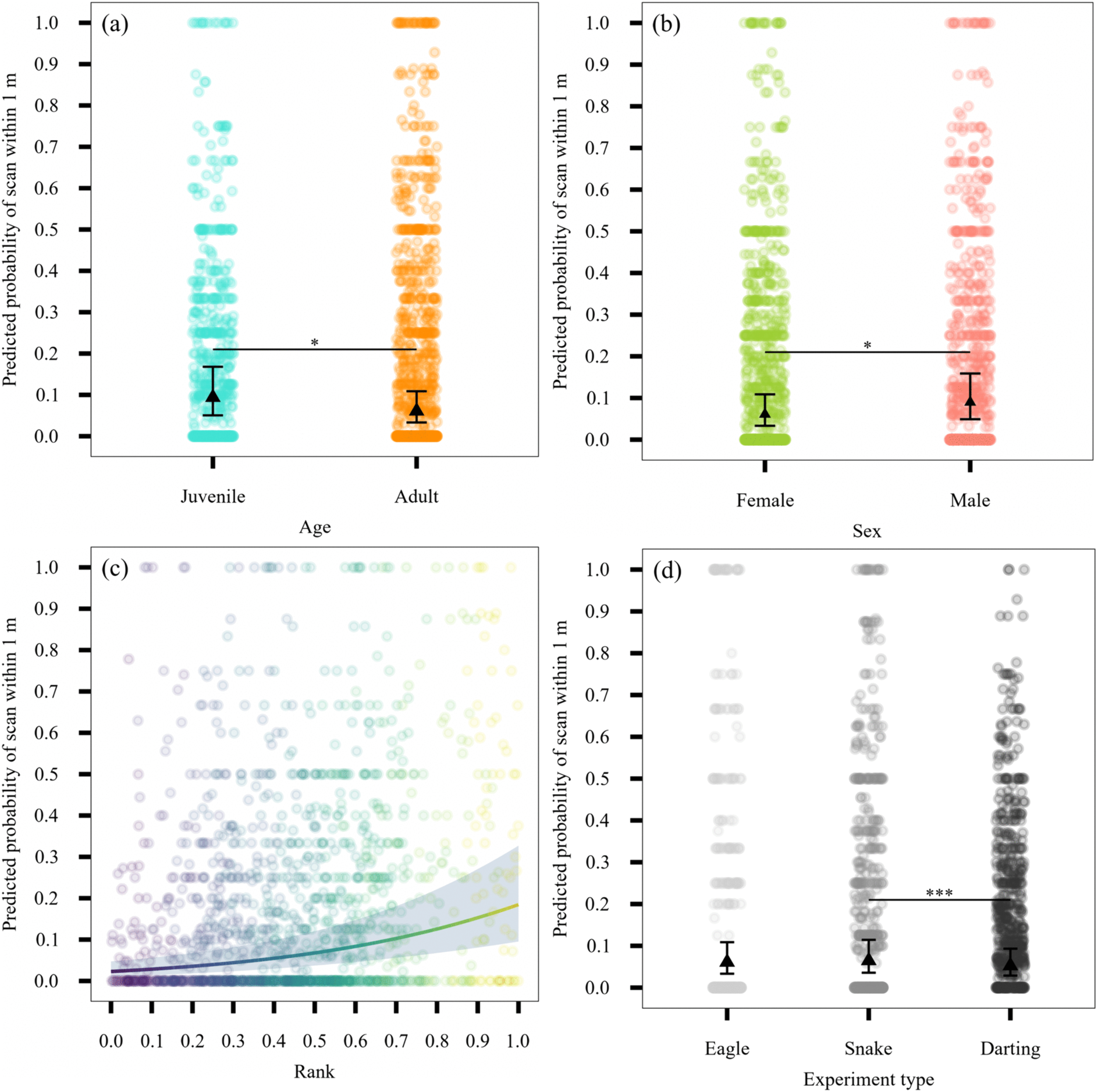
Predicted probability of an individual being within 1 m of the corn during scans across demographic and experimental variables. Panels show the effects of (a) age, (b) sex, (c) rank, and (d) experiment type. Individuals are represented as points. In panels a, b, and d, central points represent model-predicted probabilities and error bars indicate 95% confidence intervals. In panel c, the solid line represents the predicted probability, and the shaded area indicates the 95% confidence interval. Significance levels are indicated as follows: * p < 0.05; ** p < 0.01; *** p < 0.001.

In ZINB_1, age had a significant impact on number of scans spent within 1-5m of corn (LRT= 16.18; p < 0.0001) with juveniles having 1.75 times (75%) higher odds than adults to be within 1–5m of corn (Estimate = 0.56; p < 0.001). We found a significant effect of rank on the number of scans spent within 1-5m of corn (LRT = 4.00; p = 0.045) with higher-ranking individuals having 1.77 times (77%) higher odds, per unit of standardized rank, than lower-ranking individuals to be within 1–5m of corn (Estimate = 0.57; p = 0.050). Neither sex (LRT=0.74; p = 0.39) nor experimental condition (LRT=2.04; p > 0.05) significantly influenced proximity within 1 to 5m of corn. The random effect of ID nested within the group had a variance of σ² = 0.53 (SD = 0.73) and the group alone had a small variance of σ² = 0.07 (SD = 0.27). The variance from trial number was small (σ² = 0.03; SD = 0.18). The marginal R² was 0.079, indicating that the fixed effects explained only 7.9% of the variance in the amount of time spent within 1 to 5m of the corn. The conditional R² was 0.70, suggesting that the full model—including the random effects of ID nested within the group and trial number explained 70% of the variance.

#### Effects of age, sex, rank and experiment on rate of alarm calling and model approaches

In the conditional model of ZIP_1, sex had a significant effect on giving alarm calls (LRT = 6.22; p = 0.013) with male vervet monkeys alarm calling at a rate 146% higher than females (Estimate = 0.90; SE = 0.30; z =3.27; p = 0.003, Fig 3a). Experiment type also had a significant effect on alarm calls (LRT =47.49; p <0.0001); Tukey post hoc contrasts showed that individuals gave93% lower rate of alarm calls in the darting condition compared to the eagle condition (Estimate = -2.66; SE = 0.53; z = -5.06; p < 0.0001; Fig 3b) and 94% lower rate in the darting condition compared to snake (Estimate = -2.78; SE = 0.50; z = -5.61; p < 0.0001). There was no significant difference between snake and eagle (Estimate = 0.12; SE=0.18; z = 0.71; p > 0.05). Age (LRT = 0.30; p > 0.05) and rank (LRT = 0.36; p > 0.05) had no significant effects on the number of alarm calls given.

**Fig 3.**
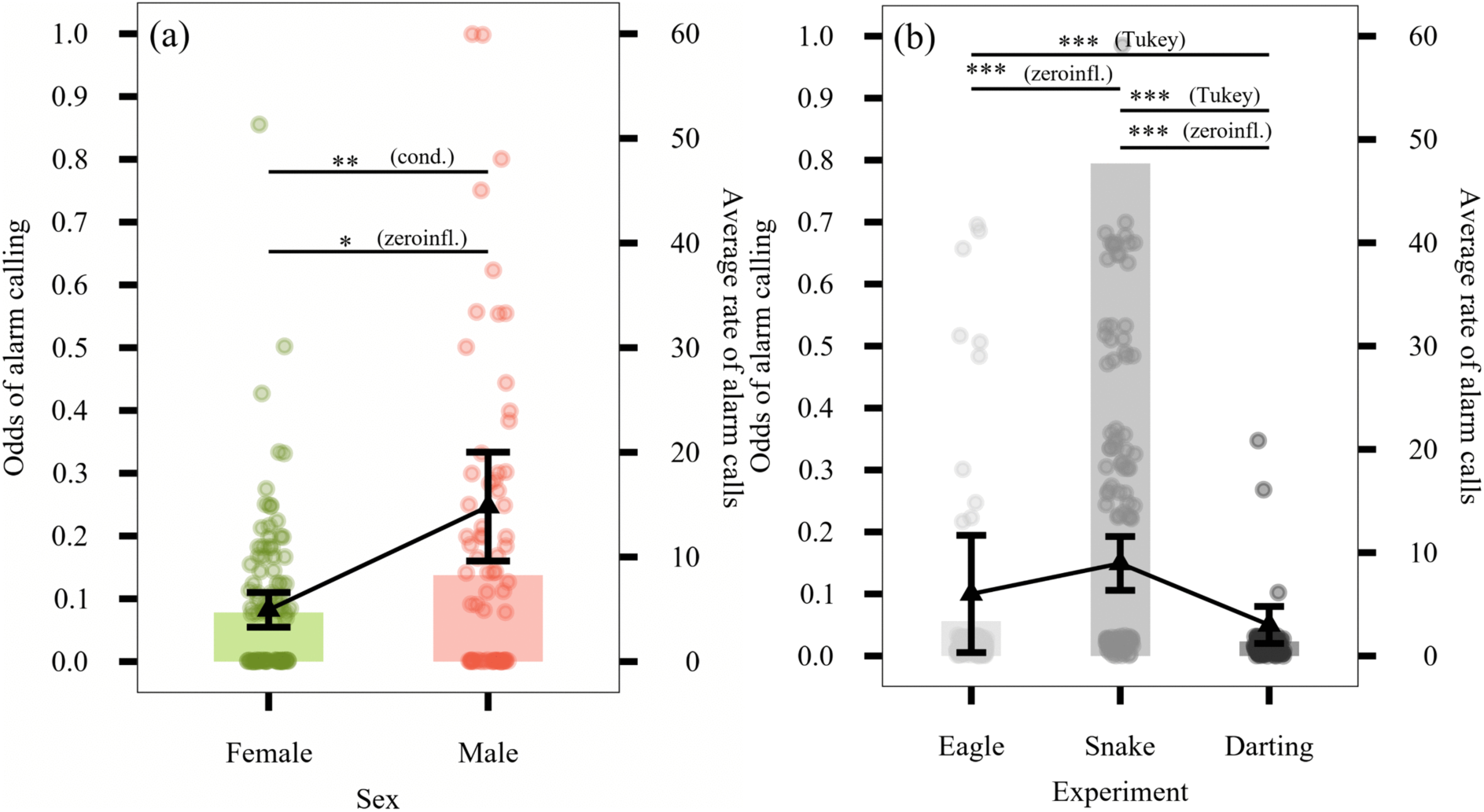
Effects of sex and experiment type on alarm-calling behaviour. Panels show effects of (a) sex and (b) experiment type on the odds of alarm calling at least once during an experiment (left axis) and the average rate of alarm calling throughout an experiment (right axis). Bars show the odds of producing at least one alarm call. Each point represents the odds of alarm calling per individual (in plot (b) 29 points were above 1.0 and were therefore removed from the plot to improve visibility). The solid line and central points represent model-estimated mean alarm-calling rates among callers, with error bars denoting 95% confidence intervals around this mean. Significance levels are indicated as follows: * p < 0.05; ** p < 0.01; *** p < 0.001. “Cond.” denotes effects from the conditional model component and “zeroinfl.” from the zero-inflation component; Tukey indicates post hoc pairwise comparisons.

In the zero-inflated model of ZIP_1, males have 40% lower odds of being structural zeros (not alarm calling), meaning that males were more likely to give alarm calls compared to females (Estimate = -0.52; SE = 0.26; z = -1.97; p = 0.049). High-ranking individuals had 67% lower odds of being structural zeros, meaning they were more likely to give alarm calls compared to low-ranking individuals (Estimate = -1.11; SE = 0.47; z = -2.38; p = 0.02). Post-hoc Tukey-adjusted comparisons of the zero-inflation component revealed that individuals had significantly higher odds of being structural zeros in the eagle condition compared to the snake condition (Estimate = 2.56; SE = 0.32; z = 7.97; p < 0.0001), corresponding to an approximately 92% increased odds of alarm calling in snake trials. Individuals also had higher odds of being structural zeros in the darting condition compared to the snake condition (estimate = 3.65; SE = 0.59; z = 6.18; p < 0.0001), corresponding to an approximately 97% increase in the odds of alarm calling in snake trials. There was no significant difference between the eagle and darting conditions (p > 0.05). Together, these results indicate that individuals were more likely to produce alarm calls during snake encounters than during eagle or darting conditions. There was no significant effect of age (p >0.05).

There was significant variance in the random effect ID nested within group (σ² = 1.48; SD = 1.22), with additional variance at the group level (σ² = 0.50; SD = 0.7) but little in trial number (σ² = 0.07; SD = 0.27). The marginal R² was 0.21, indicating that the fixed effects explained around 21% of the variance in frequency of alarm calling. The conditional R² was 0.44, suggesting that the full model—including the random effects of ID nested within the group and trial number—explained 44% of the variance.

In GLMM_2, age had a significant effect (LRT = 4.49; p = 0.034) with juveniles having a significantly higher approach rate (83%) than adults (Estimate = 0.61; SE = 0.29; z = 2.08; p = 0.038; Fig 4a). Sex and rank did not significantly improve model fit (Sex: LRT = 0.56; p = 0.456; Rank: LRT = 0.001; p > 0.05) with no differences between sexes (Estimate = 0.188; SE = 0.25; z = 0.75; p > 0.05) or across rank (Estimate = 0.02; SE = 0.56; z = 0.03; p > 0.05). Experiment type had a significant effect on the rate of approaching the model (LRT = 140.43; p < 0.0001), Tukey post hoc contrasts showed that individuals had significantly (15.2 times) higher approach rates during the snake compared to the eagle experiment (Estimate = 2.72; SE = 0.72; z = 3.79; p < 0.0001; Fig 4b) and (21 times) higher approach rates in the snake compared to the darting (Estimate= -3.03; SE = 0.33; z = -9.25; p < 0.0001). There was no significant difference between darting and eagle (Estimate = -0.31; SE = 0.77; z = -0.40; p > 0.05). There was a smaller effect of ID nested within group (σ² = 0.44; SD = 0.67) than at the group level (σ² = 1.96; SD = 1.40). There was some variance across trial number (σ² = 0.38; SD = 0.62). The marginal R² was 0.14, indicating that the fixed effects explained around 14% of the variance in model approaches. The conditional R² was 0.34, suggesting that the full model—including the random effects of ID nested within group and trial number—explained 34% of the variance.

**Fig 4.**
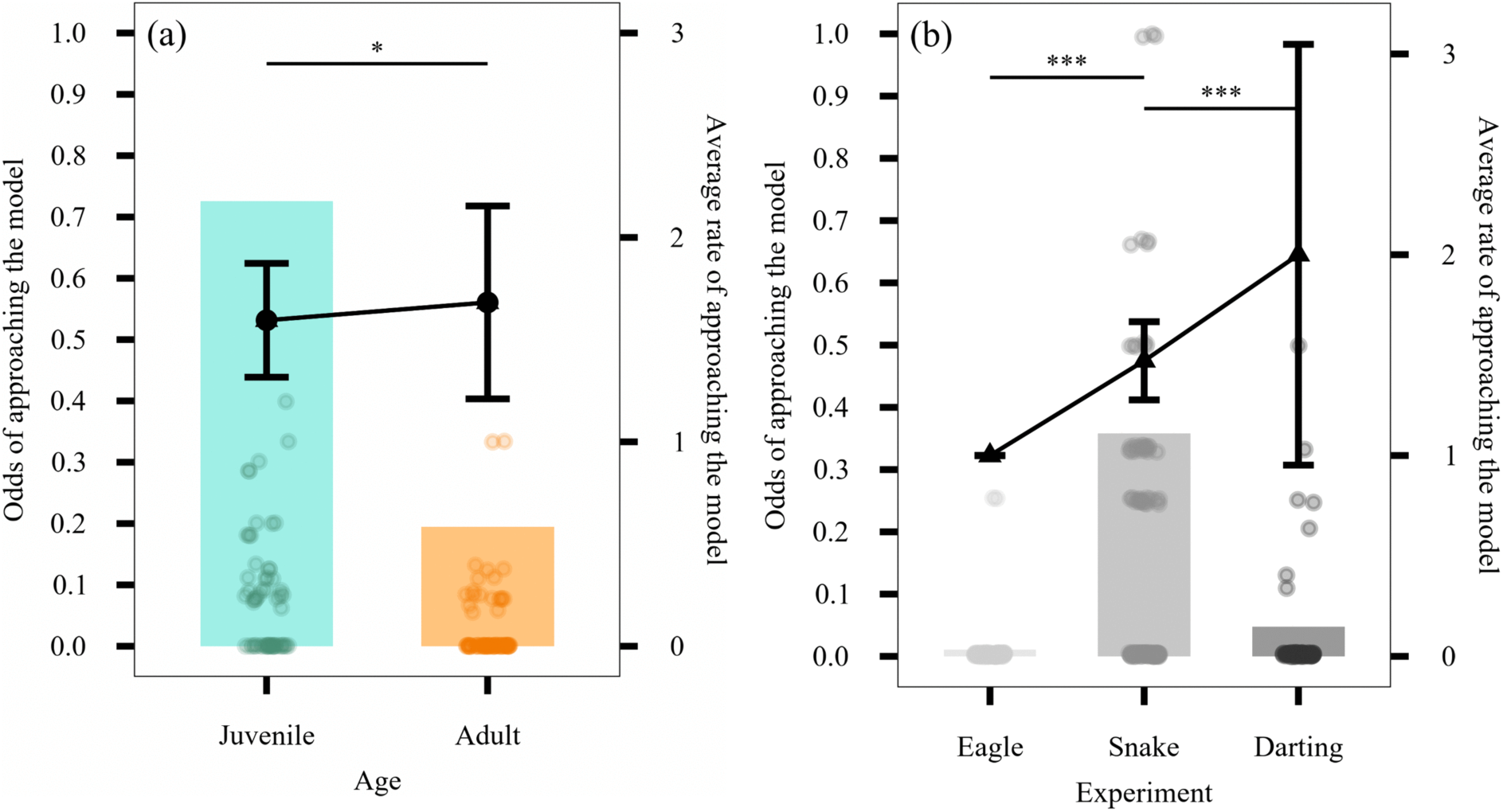
Effects of age and experiment type on approaches to the model. Panels show the effects of (a) age and (b) experiment type on the odds of approaching the model at least once during an experiment (left axis) and the average rate of approaching the model throughout an experiment (right axis). Bars show the odds of approaching the model at least once during an experiment. Each point represents the odds of approach per individual (in plot (b) 2 points were above 1.0 and were therefore removed from the plot to improve visibility). The solid line and central point represent model-estimated mean approaches rates, with error bars denoting 95% confidence intervals (CIs) around this mean. For the eagle experiment, the mean, CI low and CI high are all equal to one. Significance levels are indicated as follows: * p < 0.05; ** p < 0.01; *** p < 0.001. “Cond.” denotes effects from the conditional model component and “zeroinfl.” from the zero-inflation component; Tukey indicates post hoc pairwise comparisons.

#### Effects of sex and rank on testosterone and cortisol

In LM_1 (S1 Table), we found no significant effect of sex on testosterone levels (RSS = 243.60; p > 0.05; Fig 5a) and there was no significant difference between sexes (Estimate = 1.67; SE = 1.01; t = 1.65; p > 0.05), although there was a trend towards males having higher levels of testosterone. Rank did not improve model fit (RSS = 226.28; p > 0.05) and there were no differences across rank (Estimate = -0.65; SE = 1.48; t = -0.44; p > 0.05). The adjusted R² was 0.023 meaning our model explained around 2.3% of the variance in the data.

**Fig 5.**
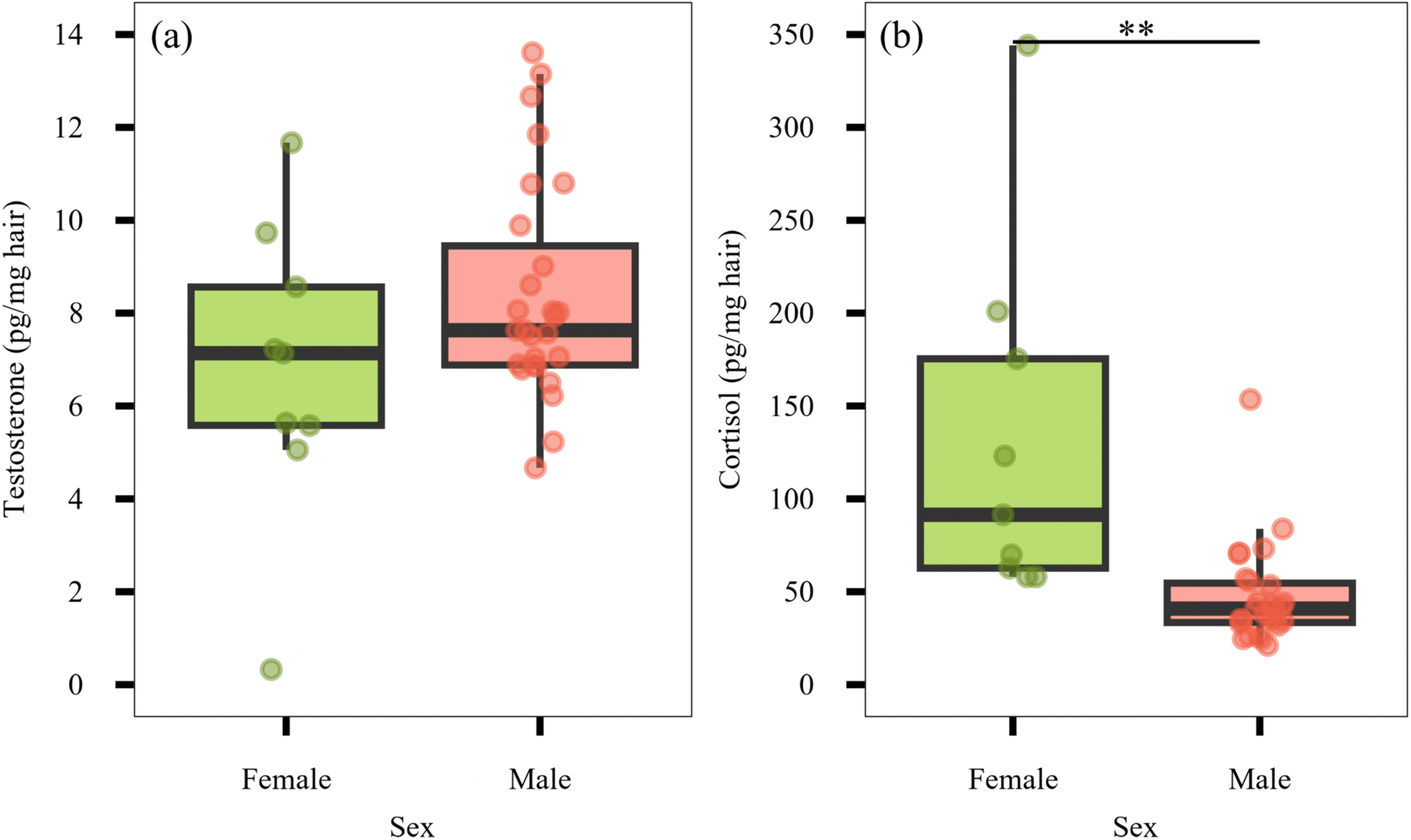
Sex differences in hormone concentrations. Concentrations of (a) testosterone, (b) cortisol (pg/mg hair) by sex. Boxes represent the interquartile range (IQR) with the median shown as the central line; whiskers extend to 1.5 × IQR. Each point represents individual concentration values. Significance levels are indicated as follows: * p < 0.05; ** p < 0.01; *** p < 0.001.

In LM_2 (S1 Table), the addition of sex improved model fit (RSS = 14.13; p < 0.0001) with males having lower cortisol than females (Estimate = -0.93; SE = 0.19; t = -4.82; p < 0.0001; Fig 5b). Rank did not improve model fit (RSS = 8.36; p > 0.05) with no differences across rank (Estimate = -0.15; SE = 0.29; t = -0.51; p > 0.05). The adjusted R² was 0.385 meaning our model explained around 38.5% of the variance in the data.

#### Effects of hormonal levels on behaviours

In GLMM_3 investigating how hormones influenced “scans within 1m of the corn”, there was no significant effect of cortisol (Estimate = -0.001; SE = 0.003; z = -0.29; p > 0.05) and this predictor did not improve model fit (LRT = 0.082; p > 0.05). Testosterone also had no impact on the number of scans spent within 1m of corn (Estimate = -0.04; SE = 0.08; z = - 0.531; p > 0.05) and it did not improve model fit (LRT = 0.28; p > 0.05). The addition of experiment type significantly improved model fit (LRT = 33.69, p < 0.001), with least time spent at the corn during the darting experiment (Estimate = -0.80; SE = 0.16; z = -5.02; p < 0.001). Tukey post-hoc contrasts showed that whilst there was no significant difference in scans spent within 1m of the corn between snake and eagle experiment types (Estimate = - 0.27; SE = 0.16; z = -1.63; p = 0.228), monkeys spent significantly less time within 1m of the corn in the darting (55% lower odds) compared to the eagle (Estimate = -0.80; SE = 0.16; z = -5.02; p < 0.0001) and in the darting (41% lower odds) compared to the snake (Estimate = - 0.54; SE = 0.12; z = -4.52; p < 0.0001). There was substantial variability across individuals in time spent at the corn (σ² = 1.435; SD = 1.198). The marginal R² was 0.068 meaning our fixed effects explained 6.8% of the variance in time spent within 1m of the corn, and the conditional R² was 0.853 showing that with the addition of the random effect of ID our model explained 85.3% of the variance in this behaviour.

In GLMM_4 investigating how hormones influenced the rate of alarm calls, testosterone did not improve model fit (LRT = 0; p > 0.05) and had no significant effect on the alarm calls rates (Estimate = −0.009; SE = 0.240; z = −0.04; p > 0.05). Cortisol did not improve model fit (LRT = 0.09; p > 0.05) and had no impact on the alarm calls rates (Estimate = −0.002; SE = 0.008; z = −0.31; p > 0.05). Again, there was considerable variance between individuals (σ² = 6.304; SD = 2.511), and a significant effect of experiment type (LRT = 375.50; p < 0.0001). Tukey post-hoc comparisons revealed a significantly lower rate of alarm calling in the darting (∼95.6%) compared to the eagle (Estimate = -3.13; SE = 1.29; z = -2.42; p = 0.038) and darting (∼99.84%) compared to the snake experiments (Estimate = -6.46; SE = 1.09; z = -5.95; p < 0.001), and there was also significantly more alarm calling in the snake (∼27.7 times) compared to the eagle experiments (Estimate = 3.32; SE = 0.71; z = 4.66; p < 0.001). The marginal R² was 0.382 meaning our fixed effects explained 38% of the variance in rates of alarm calls, and the conditional R² was 0.702 meaning that with the addition of the random effect of ID our model explained 70% of the variance.

## Discussion

The objectives of the present study were two-fold: i) to test for individual consistency in behavioural responses during risky foraging situations in seven groups of wild vervet monkeys and ii) to determine whether boldness expressed in risky situations was linked with demographic and hormonal profiles of vervet monkeys. We conducted two ecologically valid model predator experiments in the field, combined with opportunistic behavioural observations and biological sampling during capture sessions. First, we extracted several principal components in each experiment type, of which the component explaining the largest proportion of variance reflected a trait-like dimension of “boldness-shyness towards the corn”. Significant differences between vervet monkey groups were observed in their average scores for this trait, suggesting a possible social effect (e.g. social facilitation) on risk-taking behaviour. Second, individuals were consistent in a range of their behavioural responses across trials and experiment types, displaying stable personality profiles. Third, although we reported significant effects of socio-demographic factors such as sex, rank and age on risk-taking behaviour, we found almost no effect of hormones on behaviour. Our findings demonstrate that both predator models and opportunistic observations during capture sessions are suitable methods for investigating personality in vervet monkeys, and combined with hormonal data, provide valuable insights into the individual and social drivers of risk-taking behaviour in a wild-living primate species.

### Risk-taking as a repeatable trait in vervet monkeys

We found that several behaviours, including “time spent at the corn”, “time spent within 1 to 5m of the corn”, and “alarm call”, were repeatable both across trials and experimental types. These findings indicate consistent inter-individual differences in risk-taking behaviour (i.e., consistency in approaching a food source over time and across different risky situations) and antipredator responses (i.e., alarm calling), supporting the presence of stable personality variation in vervet monkeys [70,71]. Such behavioural repeatability is widely recognised as a key indicator of personality in the literature [17,19,72]. These findings provide a valuable contribution to our understanding of the behaviour and ecology of vervet monkeys and the role of personality traits as key drivers of ecological and social processes [8,17].

Repeatability estimates (r) varied across experiment types, with the highest values observed during snake and eagle model trials for both “time spent at the corn” and “time spent within 1 to 5 m of the corn”, and the lowest during darting trials. In addition, individuals spent significantly more time within 1m of corn during the snake experiment and emitted more alarm calls during both model predator experiments compared to the darting experiment.

Together, these results support the interpretation that higher perceived predation risk is associated with greater behavioural consistency, in line with those of a field study on free-ranging grey mouse lemurs by Dammhahn and Almeling [4], who simulated low and high predation risks at artificial feeding sites. The authors found that consistent individual differences in risk-taking behaviour were higher under high predation risk (i.e. when the feeding platform was on the ground) compared to low predation risk (i.e. when the platform was 1.5m high) [4].

Compared to the risks associated with predators, darting represents a more unfamiliar, less visible and less frequently encountered threat. Individuals may thus have been uncertain how to respond and may not yet have developed consistent behavioural tendencies in this context, resulting in lower repeatability. The vervet monkeys we study experience contradictory interactions with humans: they are daily followed by field assistants and researchers yet also may be chased by poachers and villagers [73]. Nevertheless, the observed reduction in time spent foraging during darting trials, suggests that this context was perceived as potentially threatening. Additionally, because sedation and capture were also carried out in the previous year using identical methods, the monkeys may have remembered this event and associated it with increased risk. Few studies have examined behavioural and physiological responses to capture and handling of wild species, and they have generally reported only short-term impacts. For example, some species show avoidance of capture equipment such as cages, but no other detectable changes in behaviours such as space use, vigilance, or foraging following capture (e.g. baboons (*Papio hamadryas*) and vervet monkeys: [74]; Samango monkeys (*Cercopithecus albogularis schwarzi*): [64]). Red colobus monkeys (*Procolobus rufomitratus*) respond to darting and collaring in a manner comparable to their responses to predatory attacks by chimpanzees, with short-term spikes in cortisol signalling an acute stress response but levels not remaining elevated for long [65].

Individuals demonstrated appropriate responses to the different experiment types, with most “alarm calls” and “model approaches” occurring during snake trials compared to eagle and darting trials. This was expected as during naturalistic encounters with snakes, vervet monkeys display specific alarm calls and a ‘mobbing’ behaviour which is hypothesised to serve to alert group members to the danger and recruit them in vigilance [75–77]. Similar responses have been observed in previous research using snake models [59,77]. In contrast, eagle encounters are characterised by a predator-specific alarm call, upward vigilance and rapid retreat into dense vegetation [60], which aligns with our finding of fewer model approaches and increased flight to nearby trees during eagle trials.

The “alarm call” behaviour showed two notable patterns: while significantly more individuals called during snake than during eagle experiment types, those that did call showed similar rates across both snake and eagle experiment types. However, repeatability in alarm calling was higher in eagle trials than in snake trials. This again likely reflects differences in antipredator strategies, with snake encounters eliciting more variable, context-dependent responses, and aerial predators requiring more rapid and consistent signalling. While raptors pose a high-risk and rapidly unfolding threat, snakes are less mobile, and primate responses to them seem to be more variable and context-dependent, varying with factors such as the snake’s location [78] and species [77].

During our control turtle trials, individuals spent significantly more time at the corn and rarely produced alarm calls. This indicates that the stimulus was not perceived as a predator or likely threat (S1a-c Fig). However, rates of vigilance were higher in the turtle and snake compared to the eagle and darting, suggesting that individuals were still cautious of this novel object and use vigilance to scan the grass. Similar responses to novel objects have been reported previously in vervet monkeys [23]. This result further supports the validity of the predator model experiments, as behavioural responses were predator-specific and consistent with natural antipredator strategies. Taken together, these findings demonstrate that risk-taking and antipredator behaviour in vervet monkeys are repeatable traits expressed across ecological contexts, with the degree of consistency shaped by the perceived level and type of threat.

### Group-level differences in boldness

Our analyses revealed clear group-level differences in boldness towards the corn. Individuals from AK and KB groups consistently displayed higher levels of boldness towards the corn than those from BD and LT, while no significant differences emerged among the remaining groups. These findings suggest that boldness towards the corn is not evenly distributed across the population but instead exhibits group-specific patterns, with some groups characterised by greater risk-taking than others. A recent study on the same population found that group-level differences in sociality were stable over time with AK being more social than BD, even after controlling for demographic factors such as sex ratio, age structure and group size, and without apparent genetic or ecological explanation [34]. Another study [79] reported that AK exhibited higher levels of co-feeding tolerance than KB in a co-feeding task. Taken together, these results suggest that vervet monkey groups may be characterised by distinct behavioural signatures, which may be maintained by mechanisms like behavioural mimicry leading to social conformity and group-specific social traditions. Although we could not account for genetic similarity between individuals within the tested groups, previous research found that group-level differences in boldness and exploration between four groups of marmosets occurred independently of genetic relatedness [36]. According to the authors, group-typical personality in marmosets was most likely a by-product of social learning: if adopting conspecifics’ behavioural style has no particular costs, it may result in behavioural convergence. Distinctions have been made between behavioural traits shaped primarily by genetic inheritance [80], long-term social learning (i.e., developmental effects), and short-term social learning or social facilitation acting on highly plastic traits [36,81]. Given that vervet monkeys exhibit female philopatry, with males dispersing between groups (often multiple times throughout their lives: [82]), this system presents valuable future avenues to disentangle the relative contributions of social facilitation, developmental and genetic effects on various behavioural traits.

### Effects of socio-demographic factors and experiment type on risk-taking behaviours

As supported by findings in different species (e.g. mice (*Mus musculus*): [83], grey mouse lemurs: [4]; chimpanzees: [37,84]) including humans, where risk-taking peaks in adolescence and decreases in older age [85,86], we found that juvenile vervet monkeys exhibited bolder responses during our experiments compared to adults. Juveniles spent significantly more time at the corn and within 1 to 5m of the food source than adults, and approached the predator models more frequently than adults, reflecting higher risk-taking tendencies (e.g. [87]). This is in accordance with previous studies in rhesus macaques (*Macaca mulatta*) and vervet monkeys in which young individuals were more exploratory towards novel objects than adults [23,42,66,88,89]. In human adolescents, heightened risk-taking and impulsivity have been associated with pubertal hormones, which are hypothesised to play a key role in regulating emotional processes and decision-making [90].

With regards to sex, we found that males spent more time at the corn than females, suggesting higher risk-taking, and aligning with previous findings in several animal species reporting higher boldness scores in males (chimpanzees and humans: [37]; rats (*Rattus norvegicus*): [91], Tibetan macaques (*Macaca thibetana*): [92]; crested macaques (*Macaca nigra*): [93]. These patterns are broadly consistent with life-history and sexual-selection theories predicting greater male risk-taking in humans (e.g. [94]). A recent finding on the same vervet monkey population found that males led group progression across two types of high-risk terrains, however found no direct mating advantages to this behaviour [95]. This finding supports the idea that males might be more risk-prone, but interestingly the absence of clear mating benefits raises questions as to the selective pressures driving this pattern. Indeed, a meta-analysis indicates that sex differences in personality traits are context-dependent rather than universal [96], highlighting the importance of ecological and social factors in shaping behavioural expression.

Interestingly, we found no effect of sex on the likelihood of approaching the predator models or the darting hide. This is in contrast to Schad and colleagues’ study [77] who found that adult male vervet monkeys were the least likely age-sex class to approach a snake during natural encounters. In their study, this pattern was possibly attributed to adult males often being solitary and peripheral to the group and therefore less likely to be in proximity to a detected predator. In our experiments, however, adult males were typically present during our experiments (likely because of the presence of food), which could explain why they were equally likely as females to be seen approaching the model. Our model structure further suggested that group-level factors played an important role: the random effect of ID alone was very small compared to identity nested within group, suggesting that there may have been strong differences in the tendency to approach models between groups, which could reflect socially facilitated responses to predators [36]. Overall, both fixed and random effects explained a relatively modest proportion of the variance (34%), likely reflecting the low frequency of approach behaviour and the influence of unmeasured ecological or social factors.

Males in our study were significantly more likely than females to give alarm calls, and when calling, did so at a higher rate. This finding contrasts with expectations based on kin selection theory, which predicts higher alarm calling in females due to female philopatry and greater relatedness within groups [97], and with previous work finding no sex-based differences in alarm call probability [77]. Whilst the study by Schad and colleagues [77] focused on responses to real and model snakes in the same populations of vervet monkeys as our study, they looked at a variety of different species (each with differing threats) and their research was not done within a foraging context, which might explain the discrepancy in our results. Indeed, multiple factors may be influencing when and how much males will benefit from alarm calling. For example, they are more likely to give alarm calls when in the presence of females (i.e., potential mates) than when they are in the presence of other males [97], presumably because male acceptance into social groups is (likely) dependent on the services they provide to females such as predator defence [98]. The higher rate of alarm calling fits with some previous findings that males are more vigilant than females [59,98,99]. However, neither of these studies found higher rates of alarm calls associated with the increased vigilance. Further work would benefit from further understanding the role of male services in different contexts and how this influences male acceptance within new groups and/or their reproductive success.

Our study found that higher ranking individuals spent more time at the corn and within 1 to 5m of corn than low ranking individuals. This is likely because ranking high in dominance provides fitness-related benefits, such as preferential access to food and mating partners in primate societies [100] and it is also linked to physiological differences, including context-dependent variation in cortisol levels [67,101]. However, rank alone cannot explain all the variation in behaviour exhibited by different individuals and different groups. The impact of rank on propensity to approach the corn additionally varied between groups: specifically, AK appeared to be a socially tolerant group as most of the individuals in the group had the opportunity to come to feed at the corn whereas in other groups such as BD certain low-ranking individuals never had a chance to approach the corn. Moreover, a large proportion of the variation in “time spent at the corn” was explained by random effects (roughly 77% of the variance being explained by ID nested within Group), indicating that beyond demographic covariates, much of the behavioural variance reflects stable among-individual differences and group-level effects, consistent with previous findings that dominance does not fully account for variation in foraging behaviour [102,103].

High-ranking individuals were also more likely than low-ranking individuals to give alarm calls; however, among those that did call, the rate of calling did not differ. Previous findings reported that higher-ranking individuals produce more alarm calls [59] and this effect does not occur because dominant individuals have more kin but instead likely because high-ranking individuals obtain stronger benefits to group living and thus may benefit more from efforts to maintain group cohesion and safety [97]. A recent study also found that higher-ranking males were more likely than low-ranking males to produce ‘barks’ [104], which are frequently used both as a predator alarm call and during aggressive interactions [59]. In contrast, rank did not influence the “approach the model” behaviour.

### Effects of sociodemographic factors on hormonal levels and of hormonal level on risk-related behaviours

Regarding the effects of sociodemographic factors on hormonal levels, males exhibited significantly lower levels of cortisol than females. This is in accordance with previous studies reporting that males displayed lower cortisol levels compared to females in various primate species (common marmosets: [105]; vervet monkeys: [55,56]). In vervet monkeys, lower cortisol has also been associated with increased boldness (vervet monkeys: [55]; but also *see review* [106]). In line with this, males in our study displayed higher levels of risk-taking behaviour, suggesting that lower cortisol concentrations may be linked to behavioural tendencies such as boldness or impulsivity. Hair cortisol concentration is considered a good marker of long-term cortisol release and of chronic stress both in human and nonhuman primates [107,108], supporting the hypothesis that baseline physiological state may be associated with stable behavioural traits.

While males showed a trend towards higher levels of testosterone compared to females, this difference was non-significant. This is likely attributable to a combination of high inter-individual variation in hormone levels (e.g., adult female Praia exhibited a testosterone concentration of 11.67 whereas adult female Circe had a concentration of 0.33), and our relatively small sample size (N = 37). Our physiological data collection was opportunistic, which limited this sample size and affects the statistical power of our results. Moreover, we cannot exclude that some samples might have been degraded, either during sampling in the field, despite all the hygiene measures that have been taken, or during transport to the lab. Biological sampling in the middle of the savannah is challenging and offers a less controlled environment compared to captivity settings with laboratory and all equipment onsite. Nevertheless, we believe it was worthwhile to take advantage of this opportunity, as personality studies often lack the combination of behavioural and physiological measures on wild animals.

Contrary to our expectations and to previous studies on primate and non-primate species [46,48,55–57], we did not find any significant correlation between hormonal levels and risk-related behaviours (i.e., time spent at corn, alarm call rate). Again, this was likely due to low statistical power and warrants further investigation. While the relationship between testosterone, cortisol and risk-taking has been explored in economic decision-making contexts in humans (e.g. [46,47]), its role in anti-predator behaviour remains largely unstudied. Our findings suggest that in predator-encounter contexts, individual differences may be a stronger predictor of risk-related behaviour than hormonal state.

Our findings provide valuable insights into risk-taking behaviour in a large number of 241 wild vervet monkeys. We have demonstrated how various factors such as age, sex, rank and, to some extent, hormones, impact risk-taking behaviour as expected. Beyond this, we observed that individuals show consistent variation in various behaviours in the context of foraging under risk of predation or capture in their natural environment. We have additionally demonstrated that groups show variety in a “boldness towards the corn” trait, with individuals in the same group being more likely to behave like others in their group and thereby supporting the idea that there may be group-norms or socially acquired norms in individual responses to risk. Taken together, these results highlight the complex interplay of individual traits, social environment, and ecological context, providing one of the first comprehensive demonstrations of how stable behavioural differences and group-level tendencies jointly shape risk-taking in a wild primate population. Nevertheless, more studies are needed to provide insights into how consistent individual traits translate into natural behavioural patterns and group dynamics, to ultimately shed light on fitness consequences of personality and social roles.

## Materials and methods

### Study site and animals

Data were collected from early October 2023 until early April 2024 at the INKAWU Vervet Project (IVP), Mawana Game Reserve (28° 00.327 S, 031° 12.348 E), KwaZulu Natal, South Africa. At the beginning of the study, subjects included 241 individuals from seven groups of wild vervet monkeys: Ankhase (AK), Baie Dankie (BD), Crossing (CR), I-Family (IF), Kubu (KB), Lemon Tree (LT) and Noha (NH). Group compositions varied across the study due to deaths, births, and male dispersals (S2 Table). Females were considered juveniles until they gave birth to their first infant; males were considered juveniles until they dispersed from their natal group (typically occurring at around four years old).

All groups were habituated to human presence allowing for observations within five metres. The process of habituation began in 2010 for AK, BD, LT, and NH, and in 2013, 2014, and 2020 for KB, CR, and IF, respectively. All individuals were recognisable through natural facial and body features. Observers were trained and tested for individual identification of subjects and assessed for inter-observer reliability in behavioural variables scored with an experienced observer before participating in data collection.

We used three different high-risk scenarios within the context of foraging: one experiment with a capture risk and two experiments with a predation risk. In each experiment, monkeys were baited with a source of appetising food (i.e. rehydrated corn kernels; energetic value: 365 kcal/100g), but under a different regularity/expectancy of risk: an irregular/less expected capture risk during darting events (Figs 6a-b and 7a), and a regular/more expected predation risk during the predator model experiments (Fig 7b-d). In addition to these experiments, we ran one control condition in each group involving a non-threatening novel object (a plastic toy turtle; Fig 8c). Darting sessions occurred for the whole month of October 2023, the control experiments occurred in October and November 2023, and predator model experiments were performed over six months, i.e. from November 2023 until April 2024. No more than one experiment was conducted per group per day.

**Fig 6.**
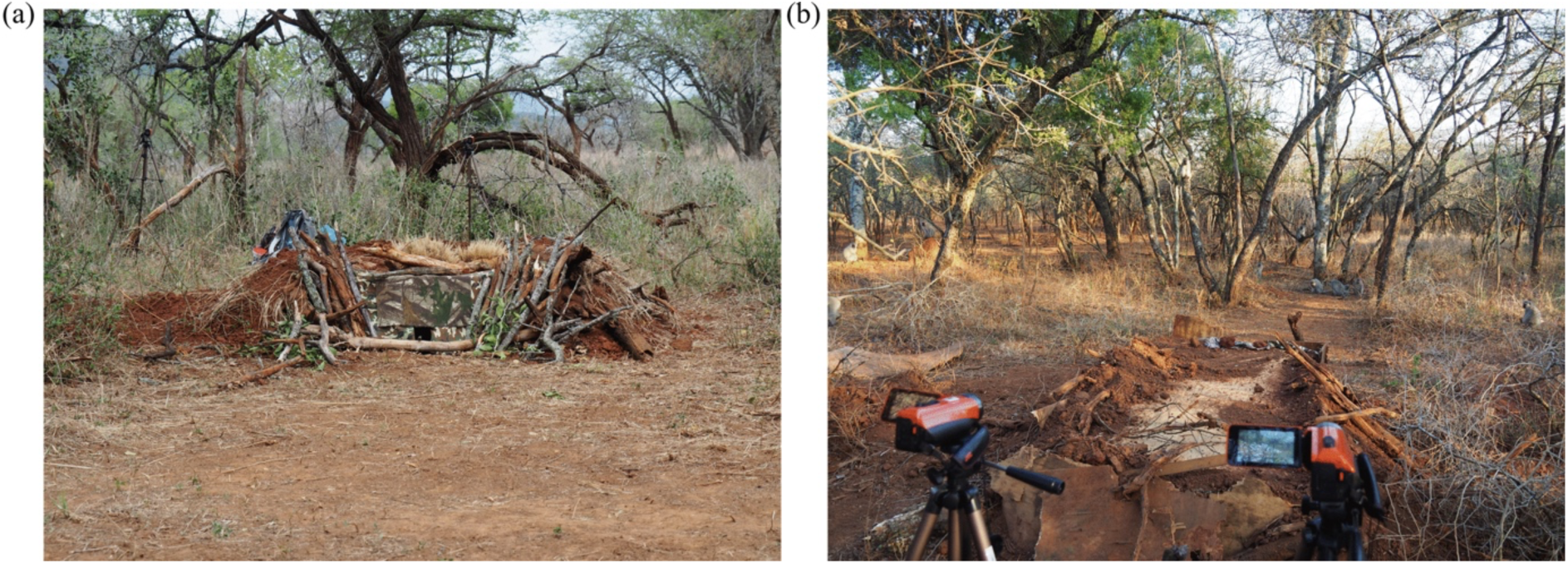
Photographs of the hides used for darting. (Copyright: Charlotte Canteloup) (a) View of the front of ‘Jacaranda hide’ (NH) from the feeding site with the camera mounted on a tripod behind the hide. The tip of the dart gun is visible through the opening. (b) View of the back of ‘Sleep 2 hide’ (BD) with two cameras mounted on tripods behind the hide. Monkeys are visible at the feeding site, 8m away from the hide.

**Fig 7.**
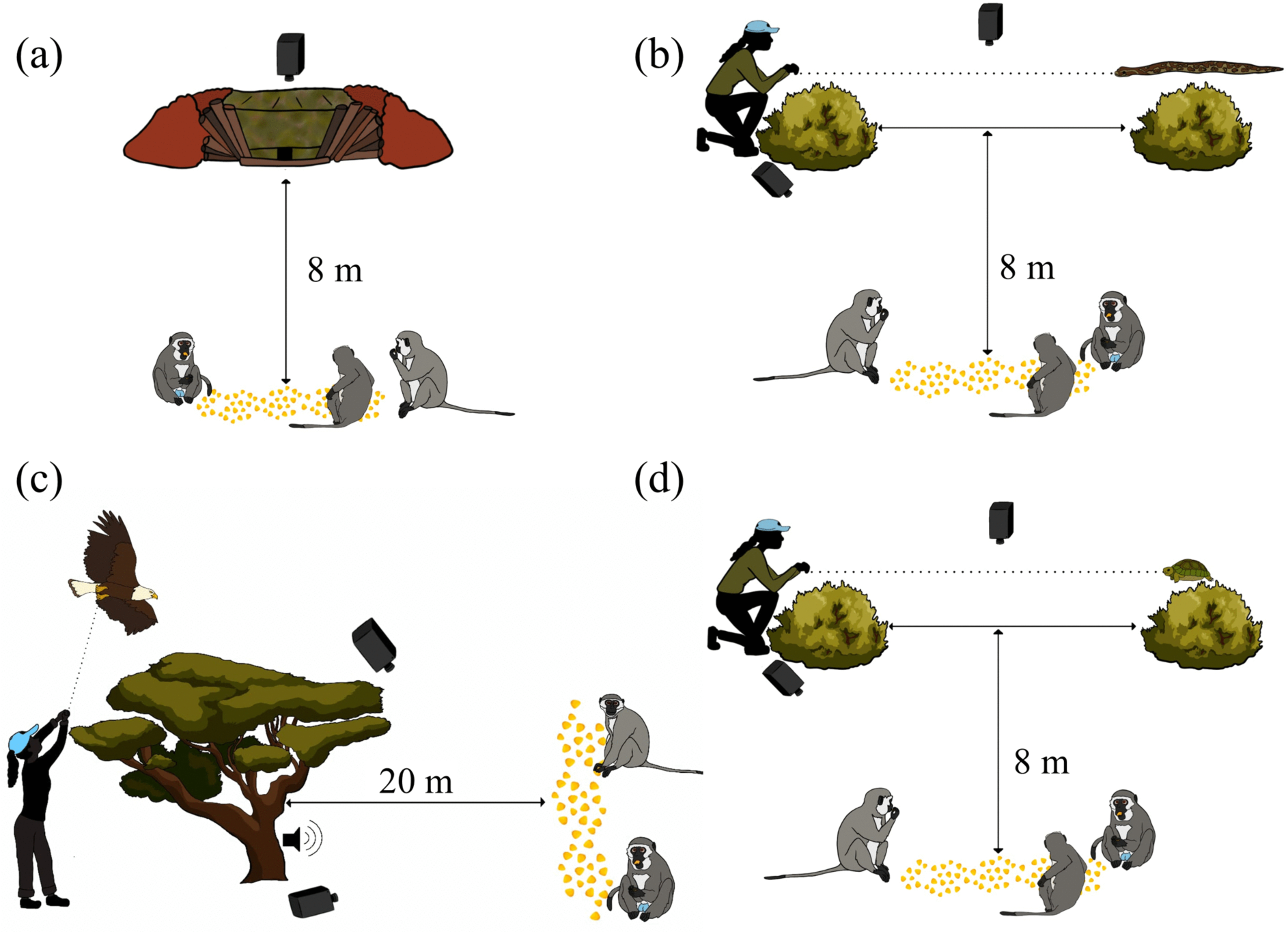
Experimental set-up for the four different types of experiment, showing the placement of the corn, camera and experimenters. **(a)** The darting experiment, **(b)** the snake model experiment, **(c)** the eagle model experiment, and **(d)** the turtle model experiment. Experimenters were behind the hide for the darting experiment, were hidden behind a large bush in the snake and turtle experiment and were behind a small tree or a bush (of an appropriate height for the kite to be visible if flown behind it) for the eagle experiment. The corn was arranged in such a way that the monkeys faced the direction of the hide or the predator. Cameras were positioned behind the hide and at a distance from the predator, affording a clear view of the monkeys and the experimental location. For the eagle experiment, the speaker emitting the eagle call was attached to a tree. (Copyright: Fanny Aguilera).

**Fig 8.**
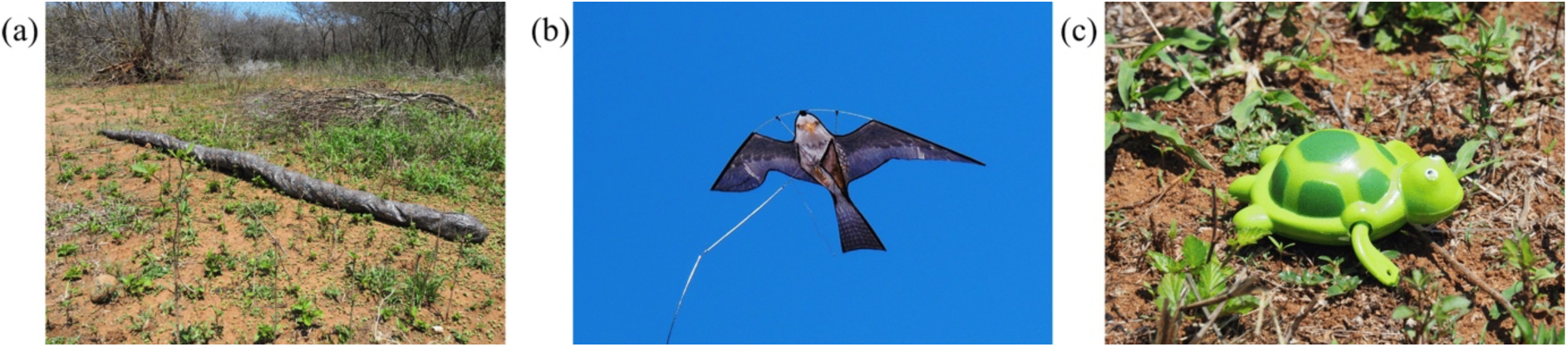
Photographs of the three different predator models used in the study. **(a)** The stuffed python (**b)** the kite eagle, **(c)** the toy turtle. (Copyright: Charlotte Canteloup).

### Experimental procedure

#### Setup

For the darting experiment, once a group was within 500 m of one of the hides at any time during the day (i.e. between 5 am and 4 pm), we went to set up the experiment at the hide. Then, the experimental session was started once the first approaching monkey arrived within one metre of the corn. Predator model experiments and the control conditions were conducted just before or as the monkeys left their sleeping sites in the morning (i.e. between 5 am and 11 am, depending on the season) to ensure that the whole group was present and able to participate in the experiment. Once the monkey group was located, the experimental setup was prepared 50-100 metres away. Then, the experimental session was started once the first approaching monkey arrived within five metres of the corn.

The setup for the experiments consisted of two cameras (Dolby JVC camcorder AVCHD) set on tripods; one was zoomed to capture a five-metre radius around the feeding site from a distance of 10 metres, and the other one was set to overview the surrounding area. The feeding site consisted in a line of corn spread along two to four metres (according to group size, e.g., two metres for KB and four metres for BD) on the ground. For the darting experiment, due to better visibility the setup with two cameras was the same when working with the largest group (BD) but included just the zoomed camera for the other six groups, and the line of corn was located eight metres away from the darting hide (Fig 7a). For the predator model experiments the line of corn was spread at either five or 20 m of the hidden predator model (Figs 7b and c). Two to five trained observers took part in the experiments by standing behind the two cameras to identify the monkeys.

Once the setup was ready, the observer at the zoomed camera continuously verbally identified all individuals feeding at the corn or present within one to five metres from the corn. The observer at the second camera identified all individuals who were visible within the surrounding area and estimated their distance from the feeding area (five to 10 metres, or outside of 10 metres). Both observers recorded key behaviours (S3 Table) such as self-scratching and alarm calling.

#### Darting: conditions of capture, anaesthesia and hair collection

Here, we took advantage of video-recordings of some capture sessions conducted for another project (see “Ethical Note” below) aiming at removing old bio-logger collars and fitting new ones to anesthetised vervet monkeys. The present project aimed to analyse monkeys’ behaviour and their propensity to take the risk to forage in this context. Darting sessions occurred in October 2023 in all seven groups, targeting 37 adult individuals (28 males; 9 females). Once anesthetised, hair samples were collected from those individuals to measure their hormonal levels. In total, 46 darting sessions took place across 24 days, starting as early as 5:00 and lasting as late as 16:45, with an average duration of 2h 02min per day (Table 1). Darting and anaesthesia monitoring were carried out by M.H., a wildlife veterinarian, expert in primate anaesthesia with extensive experience of darting wild primates in Africa and Asia (>1000 darting events). Between one and three groups were darted per day and were chosen opportunistically depending on their proximity to the hides (i.e. an underground structure used to observe animals without disturbing them). In total, 38 successful darting events took place (one individual ‘Yaz’ was darted twice as the first collar was dysfunctional). A darting trial was considered successful when it ended with the veterinarian holding an anaesthetised monkey. Some were considered unsuccessful, primarily because the target was missed. But in a few cases, the monkey ran too far away and could not be found in time; there were also two cases of injury (IF and BD).

**Table 1.**
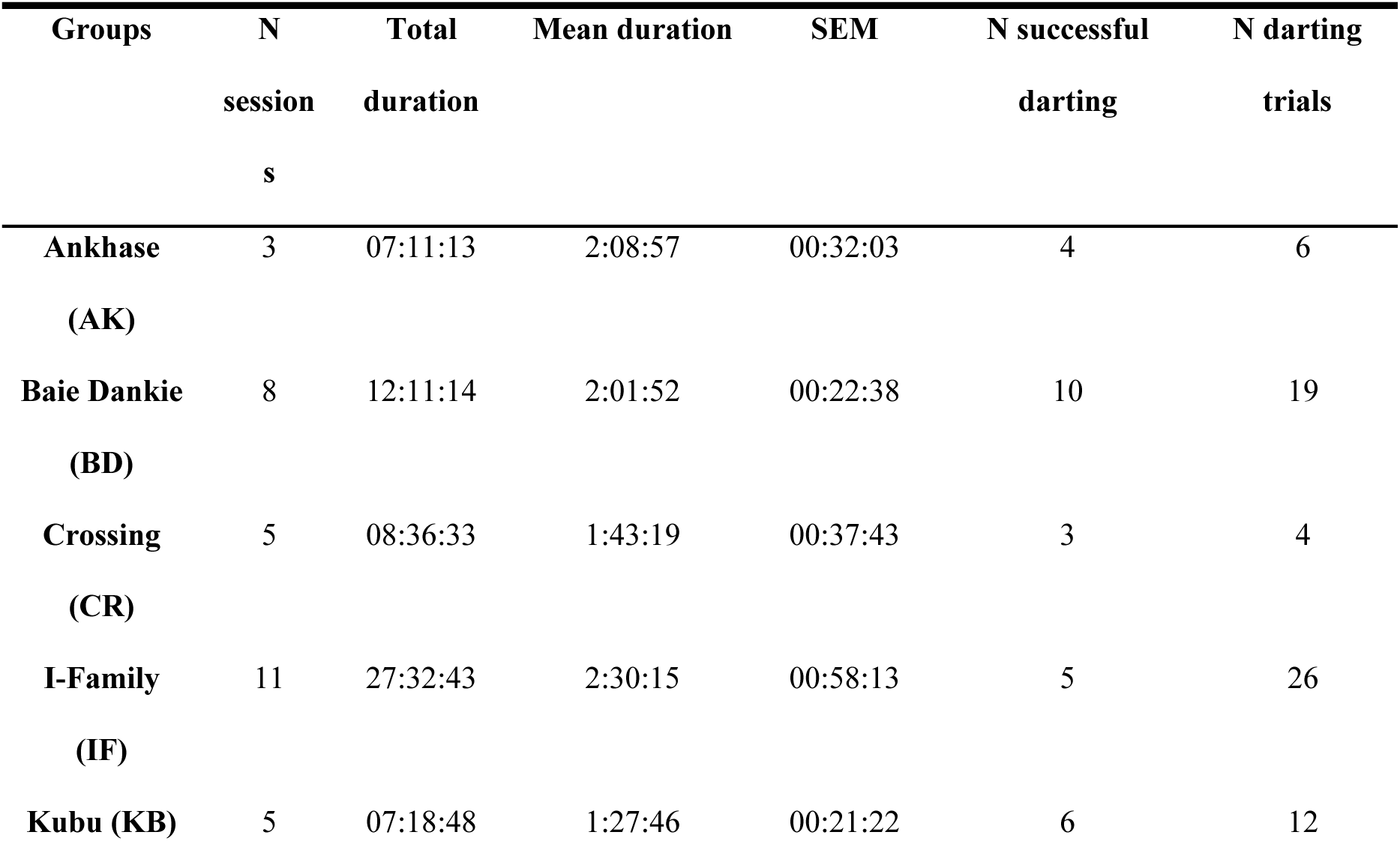
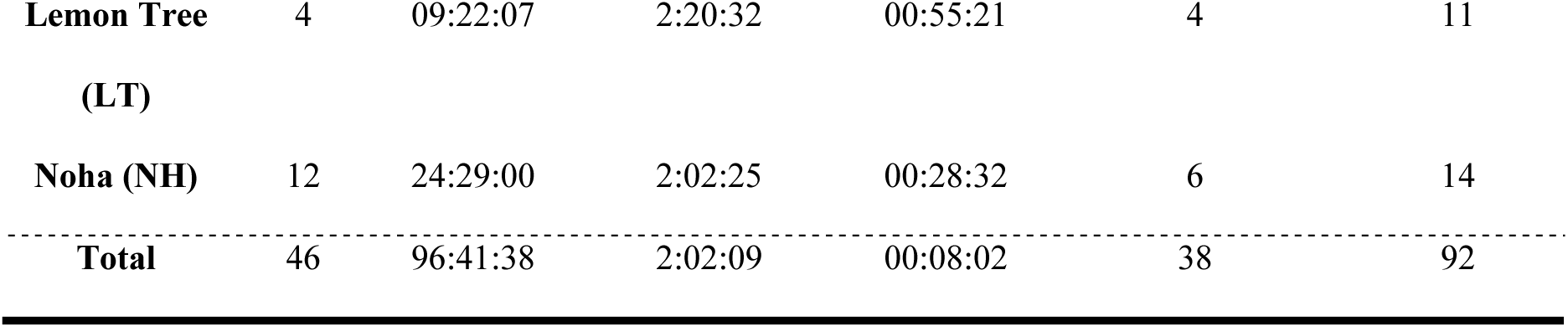
Details on the darting sessions that occurred in the seven vervet monkey groups included in this study. Total duration, mean duration and standard error of the mean (SEM) expressed in hh: mm: ss.

The capture protocol using hides has been designed by M.H to assure minimum disturbance of the animals. One to two hides per group (11 hides in total) were built in monkeys’ territories prior to the darting period. Hides were built only in patches of grassland where trees were small enough to minimize the risk of severe injuries after an eventual fall, as monkeys regularly climb trees after being darted [109]. A hide consisted of a rectangular hole (2mx1mx1m) dug in the ground, large enough for the veterinarian and his dart gun to be hidden inside (Fig 6). The hole was covered by a wooden board, some branches, foliage, and soil. A small opening (10cm x 5cm) through a camouflage blanket allowed the veterinarian to insert the dart gun and view the feeding site (Fig 6). Branches and leaves in front of the hide were pruned to avoid any obstacle that could deviate the dart from its trajectory. As soon as the veterinarian was hidden inside the hide, corn was spread in the feeding site, 8m away in front of the hide, to attract the monkeys.

Between three to five people were present during each darting session, including M.H, F.A and additional field assistants. As the sessions progressed, the corn was often monopolized by the same individuals. Therefore, we occasionally added two or three piles of corn, depending on group size, three metres away from the main pile in six of the groups (BD, CR, IF, KB, LT, NH), but still about 8m away from the hide, to increase our chances of darting all target individuals.

The veterinarian used a CO2-powered injection pistol (Daninject, Denmark) to dart the monkeys. 1.5×20 mm collared needles, mounted on 3 ml plastic darts (Teledart, Germany), were used for both males and females. Standard darts of a combination of 10 mg of tiletamine-zolazepam (Zoletil® 50, 50 mg mL-1; Virbac, France), and 0.2 mg of medetomidine (2 mg mL^-1^; V-Tech, South Africa) were used to induce anaesthesia in males. Standard darts of a combination of 6 mg of tiletamine-zolazepam (Zoletil 50, 50 mg mL^-1^; Virbac, France), and 0.12 mg of medetomidine (2 mg mL^-1^; V-Tech, South Africa) have been used to induce anaesthesia in females. The mean (+/- SD) standard induction dosages were respectively 1.98 mg kg^-1^ +/- 0.24 mg kg^-1^ for tiletamine-zolazepam and 0.04 mg kg^-1^ +/- 0.01 mg kg^-1^ for medetomidine. The veterinarian aimed at the hindquarters of the animal and tried to hit it perpendicularly. For safety reasons, vervet monkeys were darted only when located on the ground to decrease the risk of having an individual falling from a tree. When a target monkey was darted, the monkeys would flee from the feeding site. The team waited 15 min for the anaesthetic to take effect before approaching the monkey. If other monkeys returned to the feeding site during this time, another monkey could eventually be darted. If no additional monkey was darted within 15 min, the team had 30 min to find the darted monkey within a 100-meter radius around the hide. Once the monkey was found, it was collected in a blanket to be examined and processed by the veterinarian.

The anesthetized monkey was placed on a lateral recumbency for the procedure. All team members were wearing personal protective equipment (e.g. masks, gloves, long sleeves) to prevent cross species transmission of human diseases. A complete physical examination of the monkey was performed with special care at the darted hamstring muscles. Human-induced impacts of darts have been systematically cleaned with 70% ethanol and covered with aluminium in aerosol (Aluspray, Vetoquinol, France). While the monkey was anesthetised, the team weighed it, took some biological measures (e.g. teeth and testicle size) and fitted the individuals with the bio-logger collars for the needs of the other project. We collected a 4cm x 4cm hair sample from the rump on all darted individuals with an electric shaver and stored it in a paper envelope, labelled with the individual’s name and date of capture. If necessary, anaesthesia was prolonged with 1 mg kg^-1^ of tiletamine-zolazepam (Zoletil 50, 50 mg mL^-1^; Virbac, France) and 0,02 mg kg^-1^ of medetomidine (2 mg mL^-1^; V-Tech, South Africa) both injected intravenously in the saphenous vein.

At the end of the procedure, atipamezole (20 mg mL^-1^; V-Tech, South Africa) was injected in the adductor magnus muscle at an approximate medetomidine/atipamezole dose ration of 1:5. The monkey was then subsequently released near easily climbable trees within visual range of their social group and was monitored until it was awake and able to walk safely. The average recovery time of the whole anaesthesia was usually 1-2 hours. If the monkeys’ group had travelled a long distance from the hide, the monkey was brought back to its group while still anesthetised.

#### Physiological measures

Hormonal hair analyses were done by D.M and L.K. We extracted steroids from vervet hair using previously published protocol for hair-testing [110,111]. Briefly, hair was washed twice with isopropanol and left to dry overnight. Then, it was weighed and placed in a glass vial. Methanol was added and the vials were sonicated for 30 min and then incubated overnight at 50°C with gentle shaking. The methanol was collected and evaporated under a stream of nitrogen. Samples were reconstituted in 10% methanol and 90% assay diluent that was provided with the commercial enzyme-linked immunosorbent assays (ELISA) according to manufacturer’s recommendations.

Testosterone and cortisol were quantitated in hair extracts using commercial enzyme-linked immunosorbent assays (ELISA) according to the manufacturer’s recommendations (Salimetrics; Ann Arbor, MI, USA, item no. 1-3002, for cortisol, and item no. 1-2402 for testosterone). Testosterone was validated using serial dilutions of hair pool (N = 10 samples) that showed linearity between 0.5 - 20 mg hair (equivalent to 15.4 - 600 pg/ml testosterone). The serial dilutions of the pooled hair extract showed parallelism with the provided kit standards (univariate analysis of variance in SPSS; P = 0.67). The manufacturer reported antibody cross-reactivity as 36.4% with dihydrotestosterone, 21.02% with 19-nortestosterone, 1.9% with 11-hydroxytestosterone, 1.157% with androstenedione and less than 0.489% for all other steroids. Intra-assay repeatability was determined using 3 duplicates of the pool (N = 6) on the same ELISA plate. The calculated coefficient of variation was 4.04%. Recovery was calculated as 126% by spiking a known amount of exogenous testosterone.

Serial dilutions of the hair pool (N = 10 samples) for cortisol validation showed linearity between 0.5 - 50 mg hair (equivalent to 0.037 - 3 μg/dL cortisol). Dilutions of the pool were parallel to the kit standards (univariate analysis of variance in SPSS; P = 0.093). According to the manufacturer, antibody cross-reactivity was reported as 19.2% for dexamethasone, and less than 0.6% for all other steroids. Intra-assay repeatability using 3 duplicates of the pool (n = 6) on the same ELISA plate was 4.7%. Recovery was calculated as 105.19% by spiking a known amount of exogenous cortisol.

#### Predator model experiments

Prior to running the experiments, five pilot trials were run in a recently habituated group (IF) in October 2023, to determine the most appropriate experimental procedures and to ensure that the monkeys displayed expected responses to the snake, eagle, and turtle models. This pilot data was only informative and not considered in this manuscript. Once the protocol was decided, test trials began in each of the six remaining study groups. A total of ten fake predator experiments were conducted per group (five with the python model and five with the eagle) and one control test with the turtle model was run per group. Eagle and python experiments were run alternately, with a gap of two to three weeks in between each trial. Monkeys were free to participate in the experiments and, despite our efforts to ensure and record whether the entire group was present on the day of each trial, some variability in group numbers was unavoidable (e.g., due to male dispersals or groups splitting during our study period).

#### Predator model experiments: Python

The python model (snakeskin fabric filled with cloth for shape; 3.45 m in length and 27 cm in diameter; Fig 8a) was hidden behind a bush eight metres away from the feeding site. Using a fishing line, a concealed experimenter (hidden by a camouflaged blanket) dragged the model across an open area to another bush about five metres away (Fig 7b). The model was dragged parallel to the corn line at intervals of 1.5 min for six minutes, for a total of five pulls. Once the snake was hidden behind the second bush, an experimenter covered it in a camouflage blanket and quickly moved away. Occasionally the snake remained partially visible, and the monkeys continued to give alarm calls after the six min of presentation; in which case the experimenters waited until the monkeys moved away before removing the snake (average python presentation: 7 min 7 seconds (s)). The recording continued until the corn was completely consumed after the end of the presentation.

#### Predator model experiments: Eagle

The eagle model consisted of a bird-scarer kite (137 cm wingspan; Laptony brand; Fig 8b), attached to a pole with a line extension to control flight. The experimenter was hidden behind a large bush to fly the kite, around 20 m away from the corn. In the same or adjacent bush, a loudspeaker (Ultimate Ears Roll) was hidden (Fig 7c). The model presentation lasted two minutes and consisted of flights of the kite along with martial eagle calls (obtained from YouTube; and perceived at roughly 60dB) in the following sequence: three eagle calls - flight of the kite - one eagle call - three eagle calls - flight of the kite - one eagle call. The same observational procedure was used as for the python, and recording continued until the corn was completely eaten after the end of the presentation.

#### Predator model experiments: Turtle

To control for the potential effect of novelty associated with the predator models, we used a plastic toy turtle (12 cm; Fig 8c) as a non-predator model control. We presented it only once per group in an identical set up as for the python and collected equivalent behavioural data to compare responses towards a non-predatory model with our predator models (S1 and S2 Text).

#### Ethical Note

Part of the study took advantage of an invasive procedure that was carried out for another project (ERC ‘Knowledge moves’ of E.v.d.W). This involved the capture and sedation of wild monkeys to fit bio-logger collars, as well as the collection of biological samples from a total of 37 target adult individuals (28 males; 9 females). Darting was carried out by an expert veterinarian with extensive experience in darting wild primates and the greatest care was taken for this procedure, from the design and use of the hides, the choice of anaesthetics to the monitoring of the monkeys after sedation. The University of KwaZulu-Natal animal ethics board gave permission to E.v.d.W. collaborator Prof. Colleen Downs to capture, fit a telemeter and release vervet monkeys with the assistance of a veterinarian (application AREC/00005320/2023(00020574), Project title: Ecosystem health and biodiversity: The ecology, physiology, behaviour and conservation of selected southern African vertebrates). Most of the study was exclusively non-invasive and consisted of collecting behavioural data and a minimum number of field experiments involving model predators (a model snake and a bird-kite) to test whether the animals would forage from a food source placed alongside the stimuli. The animals voluntarily took part in the experiments and were never forced to approach or be in closer contact with the risk stimuli.

All the experiments and observational data collection protocols were approved by Ezemvelo KZN Wildlife, South Africa and by the van der Walt family, the owners of the Mawana game reserve where the study took place. The University of Lausanne, Switzerland, did not have an ethics committee for the study of animals in other countries but we ensured our research adhered to the ASAB/ABS Guidelines for the Use of Animals in Research. Procedures used have been conducted in compliance with animal care regulations and applicable national laws of the Republic of South Africa.

### Behavioural coding

All videos have been coded using the video coding software BORIS v.7.13.9 [112].

During the course of the experiments, and in order to facilitate subsequent individual identification and behavioural coding, assistants conducted scan sampling [113] at two-minute intervals from the beginning of the experiment. They recorded the identity, spatial position and behaviour of each individual present (scan within 1 m, within 5 m, within 10 m, and more than 10 m away from the corn).

A total of 25 behavioural variables were coded (S3 Table). These were clustered into eight distinct categories within the ethogram: distance to corn, self-directed behaviour, agonistic behaviour, affiliative behaviour, feeding behaviour, and vigilance behaviour.

We watched all trials and coded data on each individual present at the experiment one by one. We recorded firstly whether an individual had been seen at all within the vicinity of the experimental session on that day as a binary data “Yes/No”. We then continuously coded whether an individual was present at the corn or not, to extract the total duration of “time spent at the corn”. We also counted any other observed behaviour as defined in our ethogram, such as alarm calls, vigilance, self-directed behaviours, or affiliative/agonistic interactions. For the predator experiments and the control condition, we extracted 1-minute scans to record proximity of individuals to the feeding site (i.e., within one metre, within one to five metres, within five to 10 metres, or at more than 10 metres). For the darting, 2-minute scans were recorded instead due to longer trials. This data was coded using the information provided by the assistants that had narrated what was happening during the experiments, as well as observing other behaviour/events potentially missed by the assistants (i.e. because many individuals had been simultaneously present at the feeding site).

#### Darting

During video coding, we analysed 10 min before and 10 min after each darting event, extracting a total of six scans before and after each darting trial. These 10 min were not always complete in cases where a darting occurred shortly after the start of a session, or a few min after a previous darting event. In total, 21h 36min and 8s of videos were coded. F.A. analysed all the darting videos and C.d.P. and E.B. independently coded each 10% of videos to assess inter-observer reliability of F.A.’s coding using BORIS software with a one-second timeframe comparison. The Cohen kappa was almost perfect between E.B. and F.A. (Kappa = 0.89), between C.d.P. and F.A. (Kappa = 0.92), and between E.B. and C.d.P. (Kappa = 0.86).

In addition to behavioural variables, we also coded other variables such as food calls, number of corn refills, human interference, vet movement into the hide, and darting events.

#### Predator model experiments

Videos were analysed starting from the moment of the first predator model movement until two minutes after the last movement. If the snake model was still partially visible at the end of the session, individuals continued to alarm call during the additional two minutes, and there was still corn present, we extended the coding with two additional minutes from the alarm call (average coding length for the python model was 9min 7s, and for the eagle model this was 4min 20s). A total of 7h 15min and 7s of videos were coded. Videos of the predator model experiments were coded by E.B. and C.d.P. (50% each), with 20% overlap for reliability assessment on BORIS software with a one-second timeframe comparison. The Cohen kappa was almost perfect (Kappa = 0.83).

### Hierarchical rank calculation

We used the outcomes of dyadic agonistic interactions (*aggressive behaviours*: bite, grab, hit, fight, hand-on-head, chase, stare, attack, head-bob, take place, stand-up, broad side display, frustration hop, steal food, aggressive call; *victim behaviour*: retreat, flee, leave, avoid, jump aside, crawl, self-scratch, lip smack, receiving [aggressive behaviour]; S3 Table) that were collected both *ad libitum* and during focal data collection on all adults and juveniles. This data was taken from the IVP database which consists of the project’s daily observations of all seven monkey troops.

We used the R package *EloRating* [114] to determine the linear dominance hierarchies using 11 months of data from each group (1^st^ May 2023 to 1^st^ April 2024). For every agonistic interaction, a winner (i.e. individual being the most aggressive, who used the most intense aggressive behaviour) and a loser (i.e. individual showing the most submissive behaviours and/or ending the conflict by moving away from the opponent) were determined. Based on these wins and losses, the individuals were ranked within their group using the *elo.seq* function, which adjusts the ratings of individuals over time based on the expected outcome prior to the interaction and the real outcome after the interaction. Elo scores were standardized to be scaled between 0 (lowest ranker) and 1 (highest ranker), allowing for a consistent comparison of individual rankings across different groups.

### Data analysis

#### Data Preparation

All data preparation and analyses were conducted using R (v. 4.2.3; [115]) and R studio (v. 4.4.1; [116]). For all behavioural variables collected (S3 Table), we calculated either frequencies (for all count behaviours) or durations (for “time at the corn”) per individual and corrected these values by the trial duration. Additionally, we computed composite variables consisting of several variables, that happened either rarely in the dataset or were more meaningful when combined with other variables of the same category (e.g. we created the category of “self-directed behaviour” that combined the behaviours “auto-groom” and “self-scratch”). We additionally created an ‘average’ value for each behaviour type per individual per context (i.e., one value for one individual per experiment) to be used for a principal component analysis (PCA).

#### Principal Component Analysis

Three Principal Component Analyses (PCAs) with Varimax rotation were conducted, one for each experimental context (darting, snake, and eagle), all including the same input variables to increase comparability between different experimental contexts. The PCAs were aimed to reduce the number of dependent variables and select the more meaningful ones to take into further analyses. The PCA models were constructed exclusively with variables relevant to the risk-taking context (see S3 Table, categories: distance, vigilance, self-directed), while excluding social variables (see S3 Table, categories: affiliative, agonistic). Prior to the analyses, the following variables were eliminated for various reasons: “scans within 1m of the corn” due to its redundancy with “time spent at the corn”; “head-bob the model” and “yawn” due to low occurrences; and “approach the hide” as this was only relevant to the darting. The behaviours that were included in the PCAs were as follows: “time spent at the corn”, “scans within 1 to 5m of the corn”, “scans within 5 to 10m of the corn”, “scans in >10m of the corn”, “approach the corn”, “vigilance”, “self-directed behaviour”, “flee the model”, “approach the model”, “alarm call”, and “latency to approach the corn”. For each behavioural variable listed above, one average value was provided for the PCA per individual per context (making three values per individual, if that individual had been present in all three experiment types).

Kaiser-Meyer-Olkin measures showed good sampling adequacy (darting: 0.71, snake: 0.8, eagle: 0.76), and Bartlett’s Tests of Sphericity showed significant correlations between variables in all three contexts (darting: p < 0.001, snake: p < 0.001, eagle: p < 0.001), thus running a PCA was well-suitable in all three contexts. Using the *principal()* function with varimax rotation from *psych* package in R (v2.4.6.26; [117]), four components were extracted per context based on eigenvalues >1, scree plots, and Horn’s Parallel Analysis [118], overall explaining 67.6%, 76.3%, and 69.6% of variance for darting, snake, and eagle contexts respectively.

In the darting context, we obtained four principal components with Eigenvalues >1, which together explained 67.6% of variance in our data (Table 2). The first component explained 29.9% of variance and had very high positive loadings (> 0.8) of “time spent at the corn” and “approach the corn”. The second component explained 13.8% of variance and had high positive loadings (> 0.6) of “scans within 1 to 5m of the corn” and “scans within 5 to 10m of the corn”. The third component explained 12.1% of variance and had high positive loadings of “alarm call” and “approach the model”. The fourth component explained 11.8% of variance and had very high positive loading of “scans in >10m”.

**Table 2.**
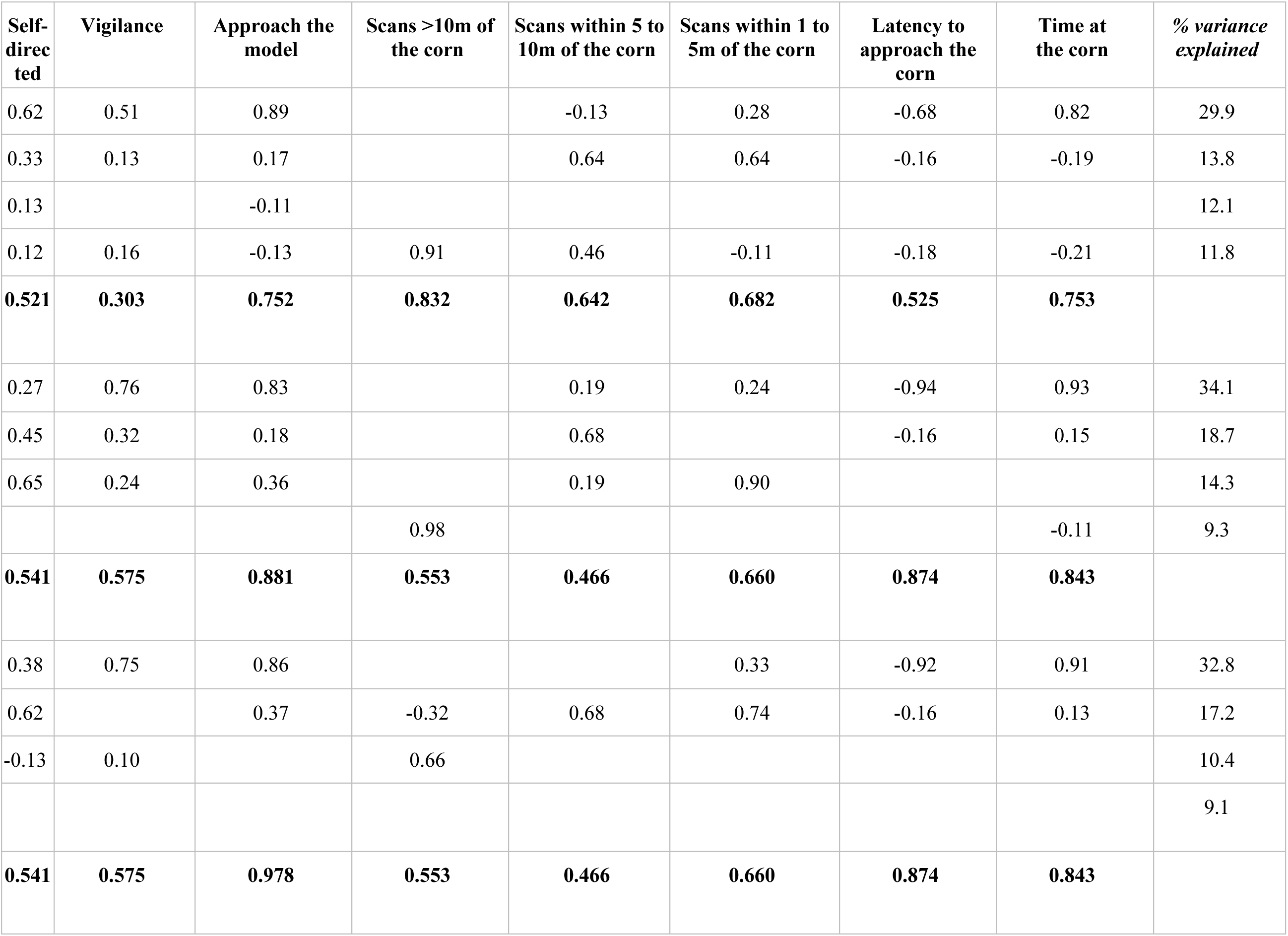
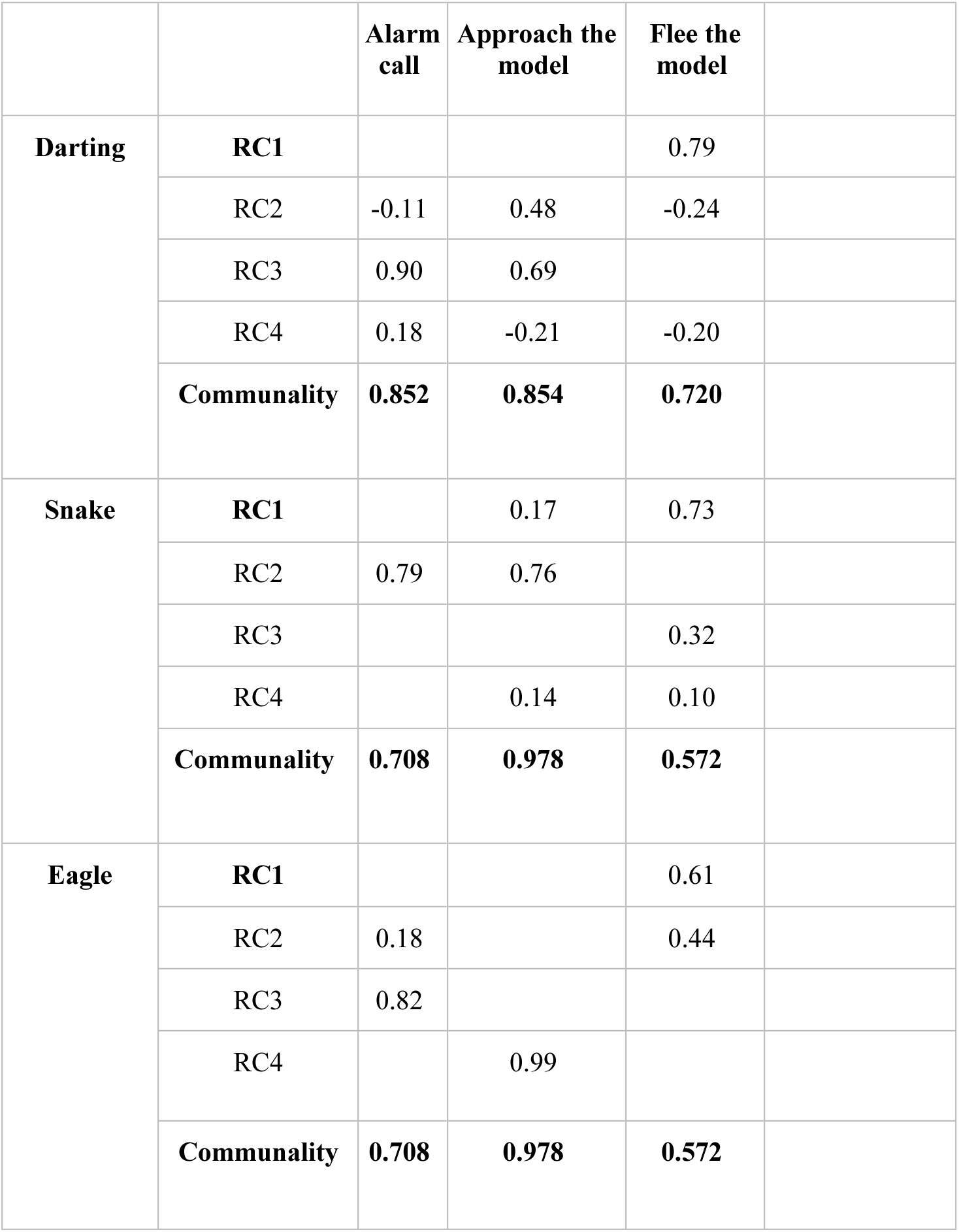
Output from the three PCAs that were conducted for each experiment type. Variable loadings for each of the main four components (PC1-PC4) per model are provided. Rows labelled PC1-PC4 represent the loading coefficients for each rotated component, and percentage variance explained by each component is provided in the final column. High positive or negative loadings indicate behaviours contributing most strongly to each component. Communality value indicates the proportion of variance in each behavioural variable explained collectively by the retained components, with higher values indicating stronger representation of that variable within the PCA model.

In the snake context, we obtained four principal components with Eigenvalues >1 which together explained 76.3% of variance (Table 2). The first component explained 34.1% of variance and had very high positive loadings of “time spent at the corn” and very high negative loadings (< -0.8) of “latency to approach the corn”. The second component explained 18.7% of variance and had high positive loadings of “alarm call” and “approach the model”. The third component explained 14.3% of variance and had a very high positive loading of “scans within 1 to 5m of the corn”. The fourth component explained 9.3% of variance and a had very high positive loading of “scans in >10m of the corn”.

In the eagle context, we obtained four principal components with Eigenvalues >1, which together explained 69.6% of variance (Table 2). The first component explained 32.8% of variance and had a high positive loading of “time spent at the corn” and a very high negative loading of “latency to approach the corn”. The second component explained 17.2% of variance and had high positive loadings of “scans within 1 to 5m of the corn” and “scans within 5 to 10m of the corn”. The third component explained 10.4% of variance and had very high positive loadings of “alarm call”. The fourth component explained 9.1% of variance and had a very high positive loading of “approach the model”.

Component 1 was consistent among the three PCAs, suggesting a trait which had loadings of behavioural variables that would best be described as bold tendencies towards the corn, and we have thus labelled “boldness towards the corn”. For darting, this component included “time spent at the corn” and “approach the corn”, and for both snake and eagle this included “time spent at the corn” and “latency to approach the corn”. We extracted individual component scores from this component to compare “boldness towards the corn” between groups. We ran a Kruskal-Wallis test to compare these individual scores between groups, and a Dunn’s test for post-hoc pairwise comparisons.

Components 1 to 4 varied a lot between experiments, therefore instead of using these components directly during our remaining statistical analyses, we decided to select and progress with the following variables in the rest of the analysis: “time spent at the corn”, “scans within 1 to 5m of the corn”, “scans in >10m of the corn”, “approach the model”, and “alarm call”. These variables were chosen as they were good representatives of each component (i.e., they loaded highly on a given component) to further investigate repeatability of behavioural responses across experiments.

#### Repeatability

To evaluate individual consistency across time and contexts, we calculated repeatability of the five behavioural variables selected from the PCAs. Due to zero-inflation and non-normal distribution of our variables, we could not use intra-class correlation coefficients, and instead fit generalised linear models using the package *rptR* (v0.9.22; [119]) to evaluate the repeatability of the binary presence/absence of each behaviour (a) across trials of the same experiment, and (b) across different contexts (experiment). For the repeatability across trials, we used the trial number as the fixed effect and individual identity (ID) as the random effect. For repeatability across contexts, we used experiment as the fixed effect, and ID as the random effect. For all models, we included 1000 bootstraps to obtain our p-values and confidence intervals (CIs). These models therefore aimed to investigate whether an individual that performed a behaviour such as “time spent at the corn”, or “alarm call”, was likely to have done so in other trials and across different experiment types as well. We did not run cross-trial repeatability for “approach the model” in eagle as there were only two occurrences of this behaviour across all of our subjects, nor for “alarm call” in darting as there were only four occurrences of this behaviour across all of our subjects. For “approach the model” in the darting experiment, we could not run bootstrapping, possibly as there were only 15 occurrences of this behaviour across all of our subjects, therefore the output of this model should be taken with caution. We did not run cross-context consistency for “scans in >10m of the corn” because it was not repeatable across-trials for snake and eagle experiments. The group IF was not included for these repeatabilities as we did not have predator experiment data for them.

#### Statistical Modelling

To test our predictions, we used generalized linear mixed-effects models (GLMMs), linear mixed-effects models (LMMs) and linear mixed models (LMs) using the packages *lme4* (v1.1-35.1; [120]) and *glmmTMB* (v.1.1.8; [121]). When we conducted pairwise comparisons using the package *multcomp*, we applied Tukey correction to the p-values. Eleven individuals had missing data for rank as they had too few data entries to obtain an accurate ELO rating and thus were removed from the models below. For each model described below, we checked for multicollinearity with the package *car* which calculates the variance inflation factor (VIF; < 1.2) (v.3.1-2; [122]) and using the package *DHARMa* (v0.4.6; [123]) we checked residuals for overdispersion, zero inflation, and outliers. If zero inflation was detected, we ran zero inflated models using the ziformula function available in glmmTMB and including the same fixed predictors as the conditional model. If there were any further issues of dispersion in such models, we then simplified the ziformula to include only an intercept. For binomial GLMMs, effect sizes are reported as odds ratios, while for Poisson-family models, effect sizes are presented as rate ratios. If there were significantly more outliers than expected (p < 0.05) we checked our data for any unusual or overly influential datapoints. A few of our models, detailed below, had a significant p-value for outliers, but as none of these appeared problematic upon inspection, and as there were always <2% of our data that were outliers, we kept these models as they were. For all models we report the main model outputs along with likelihood ratio (LRT) tests which display which predictors significantly contributed to improve model fit. R² values were obtained for each model using the package *performance* (v.0.12.0; [124]) to further evaluate model fit.

#### Comparison of “boldness towards corn” between groups

To test for differences in the first component labelled “boldness towards the corn” between groups, we used a Kruskal-Wallis test due to non-normality of the data. Following a significant overall effect, Dunn’s post hoc tests with Bonferroni correction were conducted to assess pairwise differences between groups.

#### Effects of age, sex, rank and experiment on locations relative to the corn during scans

We investigated how social and demographic factors influenced our main ‘risk’ behaviours, i.e., number of scans spent within 1m (GLMM_1; see S1 Table for the models’ details and predictions associated) and within 1m to 5m (ZINB_1; S1 Table) of the corn. We did not use the variable “time spent at the corn” because it was highly correlated with “scans within 1m of the corn” (Spearman’s rank correlation; darting: r = 0.97, p < 0.001; snake: r = 0.97, p < 0.001; eagle: r = 0.93, p < 0.001) and was zero-inflated. We did not run a model for “scans in >10m of the corn” because this behaviour did not show repeatability and because we did not have explicit predictions that we aimed to explore.

Our model investigating the effect of age, sex, rank and experiment on the number of “scans within 1m of the corn” (GLMM_1; S1 Table) was built with a binomial distribution and logit link function with “scans within 1m of the corn” as a dependent variable. For our model investigating the effect of age, sex, rank and experiment on the number of “scans within 1 to 5m of the corn”, we first ran a binomial model which displayed a significant zero inflation. Consequently, we added the ziformula with all fixed effects which created overdispersion, therefore we settled with a reduced ziformula (ziformula = ∼1) which displayed the best model diagnostics. This model (ZINB_1; S1 Table) is a zero-inflated binomial model with logit link function with “scans within 1m to 5m of corn” as a dependent variable. In both models (GLMM_1 and ZINB_1), the fixed effects included were age, sex, rank, and experiment. In both models, two random intercepts were included, first for ID nested within the group, and additionally the trial number to account for variation across trials. Post-hoc pairwise comparisons between experimental conditions were conducted using Tukey-adjusted contrasts.

#### Effects of age, sex, rank and experiment on rate of alarm calling and model approaches

To test for the factors influencing the rate of alarm calling, we fitted a zero-inflated model with Poisson distribution and log link function (ZIP_1; S1 Table), because the binomial model for “alarm call” displayed significant zero inflation. To investigate the effects of factors on the number of approaches to the model, we fitted a GLMM with Poisson distribution and log link function (GLMM_2; S1 Table). In both models (ZIP_1; GLMM_2), the fixed effects included were age, sex, rank, and experiment. Two random intercepts were included, first for ID nested within the group, and trial number to account for variation across trials. We used the count of alarm calls as the dependent variable and included trial duration time as an offset in both models to account for variation in the length of experiments. In both models, post-hoc pairwise comparisons between experimental conditions were conducted using Tukey-adjusted contrasts.

#### Effects of sex and rank on testosterone and cortisol

We created a subset of data including only individuals for whom we had hormonal data (*N* = 37). We ran two linear models (LM_1; LM_2; S1 Table) with Gaussian distribution for each hormone (testosterone, cortisol) as the dependent variable and included fixed effects for sex and rank. We did not include group as a random effect as the variance for the group was equal to 0. For the cortisol model (LM_2), we log-transformed the data due to non-normal distribution of residuals. We did not include age as a fixed effect in these models because all individuals for whom we had data were adults except one (Ndanda) who was a juvenile.

#### Effects of hormonal levels on behaviours

We ran GLMMs to test for the effect of hormones on our main behaviours of interest, “scans within 1m of the corn” and “alarm call”. We used a subset of individuals from which we had collected hormonal data. We could not run a reliable model for “approach the model” behaviour because of the combined low sample size for this behaviour and hormonal data. For “scans within 1m of the corn” we fitted a GLMM (GLMM_3; S1 Table) with a binomial distribution and ‘logit’ link function. Fixed effects included an interaction between testosterone and cortisol, as well as experiment type (i.e. to control for the variation in responses to different experimental conditions). We included only ID as random effect as the factor group contributed negligible variance and consequently resulted in singular fit. For the model investigating “alarm call”, we fitted a GLMM with a Poisson distribution and a log link function using the number of alarm calls as the dependent variable with an offset for the trial duration to account for variation in experiment durations (GLMM_4; S1 Table). We included the interaction between testosterone and cortisol, and the experiment type as fixed effects. Again, only ID was included as random effect because variance for the group was very low and this alteration improved model diagnostics. As the interaction between testosterone and cortisol was not significant (p>0.05) in GLMM_3 and GLMM_4, we removed it from the models and kept the single effects of testosterone and cortisol as fixed effects.

## Acknowledgements

We thank the INKAWU Vervet Project (Mawana Game Reserve (28° 00.327 S, 031° 12.348 E), KwaZulu Natal, South Africa) onsite managers, Mike Henshall and Siboniso Thela, and the whole IVP team for their help and support in the field, especially Nokubonga Dlamini, Zonke Mbutho, Veronicca Khosana, Matthew Monteith, Sarah Mendes, Gabriela Galindo, Josefien Tankink, Maria Granell Ruiz, Noah de Jong, Marissa Grimes, Anumita Samanta, Fabio Opreni and Pablo del Rio for their assistance during experiments. We are grateful to the van der Walt family for their permission to conduct the study on their land in the Mawana Game Reserve, KwaZulu-Natal, South Africa. We thank Lukas Schad for providing us with the python model and Christof Neumann for statistical advice. We thank the Simian Laboratory Europe (SILABE)-University of Strasbourg for hosting the team of C.C. This study has been funded by a research grant from the Fyssen Foundation granted to C.C. and by an ERC starting grant (grant agreement n°949379) granted to E.v.d.W. At the time of writing, C.C was supported by the CNRS and by NeuroStra. This work of the Interdisciplinary Thematic Institute NeuroStra, as part of the ITI 2021-2028 program of the University of Strasbourg, CNRS and Inserm, was supported by IdEx Unistra (ANR-10-IDEX-0002) under the framework of the French Program “Investments for the Future”.

## Authors contributions

Conceptualization: C.C., M.H; Data curation: C.C., E.B., C.d.P., F.A.; Formal analysis: C.d.P., F.A., C.C., V.Š.; Funding acquisition: C.C., E.v.d.W.; Investigation: C.C., E.B., C.d.P., F.A., M.H., V.Š., L.K., D.M., E.v.d.W.; Methodology field experiments: C.C., E.B., C.d.P., F.A.; Methodology darting: M.H.; Methodology IVP database: E.v.d.W.; Methodology statistical analysis: V.Š., C.C., C.d.P.; Methodology hormonal analysis: L.K., D.M.; Project administration: C.C.; Resources: C.C., E.v.d.W., L.K.; Software: C.d.P., C.C., F.A., V.Š.; Supervision: C.C.; Validation: C.d.P., C.C., V.Š.; Visualisation: F.A., C.d.P.; Writing - original draft: C.d.P., E.B., F.A., C.C., V.Š., L.K., M.H.; and Writing - review & editing: C.C., E.B., C.d.P., F.A., V.Š., M.H., E.v.d.W., L.K., D.M.

## Data availability statement

All the data and code can be found at: https://zenodo.org/records/20589623

## References

1. Fairbanks LA. Risk-Taking by Juvenile Vervet Monkeys. Behaviour. 1993;124: 57–72.

2. Pelé M, Broihanne MH, Thierry B, Call J, Dufour V. To bet or not to bet? Decision-making under risk in non-human primates. J Risk Uncertain. 2014;49: 141–166. doi:10.1007/s11166-014-9202-3

3. Arseneau-Robar TJM, Anderson KA, Vasey EN, Sicotte P, Teichroeb JA. Think Fast!: Vervet Monkeys Assess the Risk of Being Displaced by a Dominant Competitor When Making Foraging Decisions. Front Ecol Evol. 2022;10. doi:10.3389/fevo.2022.775288

4. Dammhahn M, Almeling L. Is risk taking during foraging a personality trait? A field test for cross-context consistency in boldness. Anim Behav. 2012;84: 1131–1139. doi:10.1016/j.anbehav.2012.08.014

5. Giraldeau L-A, Caraco T. Social Foraging Theory. Princeton University Press; 2000. Available: http://www.jstor.org/stable/j.ctv36zrk6

6. Lind J, Cresswell W. Determining the fitness consequences of antipredation behavior. Behav Ecol. 2005;16: 945–956. doi:10.1093/beheco/ari075

7. Janson C. Aggresive competition and individual food consumption in wild brown capuchin monkeys (Cebus apella). Behav Ecol Sociobiol. 1985;18: 125–138. doi:10.1007/BF00299041

8. Carter A, Marshall H, Heinsohn R, Cowlishaw G. Personality predicts decision making only when information is unreliable. Anim Behav. 2013;86: 633–639. doi:10.1016/j.anbehav.2013.07.009

9. Russek EM, Moran R, Liu Y, Dolan RJ, Huys QJM. Heuristics in risky decision-making relate to preferential representation of information. Nat Commun. 2024;15: 4269. doi:10.1038/s41467-024-48547-z

10. Dohmen T, Falk A, Heckman J, Schupp J, Sunde U, Wagner G. Individual Risk Attitudes: Measurement, Determinants, And Behavioral Consequences. J Eur Econ Assoc. 2011;9: 522–550. doi:10.1111/j.1542-4774.2011.01015.x

11. Roberts B, Chernyshenko O, Stark S, Goldberg L. The Structure of Conscientiousness: An Empirical Investigation Based on Seven Major Personality Questionnaires. Pers Psychol. 2005;58: 103–139. doi:10.1111/j.1744-6570.2005.00301.x

12. Farwell M, McLaughlin RL. Alternative foraging tactics and risk taking in brook charr (Salvelinus fontinalis). Behav Ecol. 2009;20: 913–921. doi:10.1093/beheco/arp059

13. Heilbronner SR, Rosati AG, Stevens JR, Hare B, Hauser MD. A fruit in the hand or two in the bush? Divergent risk preferences in chimpanzees and bonobos. Biol Lett. 2008;4: 246–249. doi:10.1098/rsbl.2008.0081

14. Montiglio P-O, Garant D, Pelletier F, Réale D. Personality differences are related to long term stress reactivity in wild eastern chipmunks (Tamias striatus). Anim Behav. 2012;84: 1071–1079. doi:10.1016/j.anbehav.2012.08.010

15. Quinn JL, Cresswell W. Personality, anti-predation behaviour and behavioural plasticity in the chaffinch Fringilla coelebs. Behaviour. 2005;142: 1377–1402. doi:10.1163/156853905774539391

16. Réale D, Gallant BY, Leblanc M, Festa-Bianchet M. Consistency of temperament in bighorn ewes and correlates with behaviour and life history. Anim Behav. 2000;60: 589–597. doi:10.1006/anbe.2000.1530

17. Réale D, Reader SM, Sol D, McDougall PT, Dingemanse NJ. Integrating animal temperament within ecology and evolution. Biol Rev. 2007;82: 291–318. doi:10.1111/j.1469-185X.2007.00010.x

18. Takola E, Krause ET, Müller C, Schielzeth H. Novelty at second glance: a critical appraisal of the novel object paradigm based on meta-analysis. Anim Behav. 2021;180: 123–142. doi:10.1016/j.anbehav.2021.07.018

19. Šlipogor V, Gunhold-de Oliveira T, Tadić Z, Massen JJM, Bugnyar T. Consistent inter-individual differences in common marmosets (Callithrix jacchus) in Boldness-Shyness, Stress-Activity, and Exploration-Avoidance. Am J Primatol. 2016;78: 961–973. doi:10.1002/ajp.22566

20. Šlipogor V, Masilkova M, Höflinger A, Lang N, Bugnyar T, Konečná M. With or without you: common marmoset, *Callithrix jacchus*, personality expression is mediated by social setting. Anim Behav. 2025;231: 123367. doi:10.1016/j.anbehav.2025.123367

21. Bosshard TC, Mundry R, Fischer J. Ecological risk-taking across age in Barbary macaques. Anim Behav. 2025;229: 123337. doi:10.1016/j.anbehav.2025.123337

22. van Dijk ESJ, Bhattacharjee D, Belli E, Massen JJM. Hand preference predicts behavioral responses to threats in Barbary macaques. Am J Primatol. 2023;85: e23499. doi:10.1002/ajp.23499

23. Blaszczyk MB. Boldness towards novel objects predicts predator inspection in wild vervet monkeys. Anim Behav. 2017;123: 91–100. doi:10.1016/j.anbehav.2016.10.017

24. Carter AJ, Marshall HH, Heinsohn R, Cowlishaw G. Evaluating animal personalities: do observer assessments and experimental tests measure the same thing? Behav Ecol Sociobiol. 2012;66: 153–160. doi:10.1007/s00265-011-1263-6

25. Stanford C. Avoiding Predators: Expectations and Evidence in Primate Antipredator Behavior. Int J Primatol. 2002;23: 741–757. doi:10.1023/A:1015572814388

26. Gonçalves A, Carvalho S. Death among primates: a critical review of non-human primate interactions towards their dead and dying. Biol Rev. 2019;94: 1502–1529.

27. Carter AJ, Marshall HH, Heinsohn R, Cowlishaw G. How not to measure boldness: novel object and antipredator responses are not the same in wild baboons. Anim Behav. 2012;84: 603–609. doi:10.1016/j.anbehav.2012.06.015

28. Isbell LA. Predation on primates: Ecological patterns and evolutionary consequences. Evol Anthropol Issues News Rev. 1994;3: 61–71.

29. Boinski S. Predators on Primates: Effects on Group Travel. Move Why Anim Travel Groups. 2000; 43.

30. Beauchamp G. Group-size effects on vigilance: a search for mechanisms. Behav Processes. 2003;63: 141–145. doi:10.1016/S0376-6357(03)00011-1

31. Rosati AG, Hare B. Decision making across social contexts: competition increases preferences for risk in chimpanzees and bonobos. Anim Behav. 2012;84: 869–879. doi:10.1016/j.anbehav.2012.07.010

32. De Petrillo F, Rosati AG. Variation in primate decision-making under uncertainty and the roots of human economic behaviour. Philos Trans R Soc B Biol Sci. 2021;376: 20190671. doi:10.1098/rstb.2019.0671

33. Šlipogor V, Massen JJM, Schiel N, Souto A, Bugnyar T. Temporal consistency and ecological validity of personality structure in common marmosets (Callithrix jacchus): A unifying field and laboratory approach. Am J Primatol. 2021;83: e23229. doi:10.1002/ajp.23229

34. Kerjean E, van de Waal E, Canteloup C. Social dynamics of vervet monkeys are dependent upon group identity. iScience. 2024;27: 108591. doi:10.1016/j.isci.2023.108591

35. van de Waal E, Borgeaud C, Whiten A. Potent Social Learning and Conformity Shape a Wild Primate’s Foraging Decisions. Science. 2013;340: 483–485. doi:10.1126/science.1232769

36. Koski SE, Burkart JM. Common marmosets show social plasticity and group-level similarity in personality. Sci Rep. 2015;5: 8878. doi:10.1038/srep08878

37. Haux LM, Engelmann JM, Arslan RC, Hertwig R, Herrmann E. Chimpanzee and Human Risk Preferences Show Key Similarities. Psychol Sci. 2023;34: 358–369. doi:10.1177/09567976221140326

38. Bunselmeyer C, Gunst N, Wandia IN, Williams RJ, Addessi E, Leca J-B. Sex and Age Differences in Decision-Making Under Risk by Wild Balinese Long-Tailed Macaques (Macaca fascicularis fascicularis): A Field Experimental Study. Animals. 2026;16: 617. doi:10.3390/ani16040617

39. De Petrillo F, Ventricelli M, Ponsi G, Addessi E. Do tufted capuchin monkeys play the odds? Flexible risk preferences in Sapajus spp. Anim Cogn. 2015;18: 119–130. doi:10.1007/s10071-014-0783-7

40. Rivière J, Kurt A, Meunier H. Choice Under Risk of Gain in Tufted Capuchin Monkeys (Sapajus apella): A Comparison With Young Children (Homo sapiens) and Mangabey Monkeys (Cercocebus torquatus torquatus). J Neurosci Psychol Econ. 2019;12: 159–168. doi:10.1037/npe0000109

41. Chaix-Eichel N, Guerillon A, Bourgeois-Gironde S, Rougier NP, Boraud T, Ballesta S. Social hierarchy influences monkeys’ risky decisions. Commun Biol. 2026;9: 578. doi:10.1038/s42003-026-09817-2

42. Santillán-Doherty AM, Cortés-Sotres J, Arenas-Rosas RV, Márquez-Arias A, Cruz C, Medellín A, et al. Novelty-seeking temperament in captive stumptail macaques (Macaca arctoides) and spider monkeys (Ateles geoffroyi). J Comp Psychol Wash DC 1983. 2010;124: 211–218. doi:10.1037/a0018267

43. Trivers R. Parental Investment and Sexual Selection. Sexual Selection and the Descent of Man. 1972. p. 378.

44. Wolf M, van Doorn GS, Leimar O, Weissing FJ. Life-history trade-offs favour the evolution of animal personalities. Nature. 2007;447: 581–584. doi:10.1038/nature05835

45. Karlsson Linnér R, Biroli P, Kong E, Meddens SFW, Wedow R, Fontana MA, et al. Genome-wide association analyses of risk tolerance and risky behaviors in over 1 million individuals identify hundreds of loci and shared genetic influences. Nat Genet. 2019;51: 245–257. doi:10.1038/s41588-018-0309-3

46. Apicella C, Dreber A, Campbell B, Gray P, Hoffman M, Little A. Testosterone and Financial Risk Preferences. Evol Hum Behav - EVOL HUM BEHAV. 2008;29. doi:10.1016/j.evolhumbehav.2008.07.001

47. Herbert J. Testosterone, Cortisol and Financial Risk-Taking. Front Behav Neurosci. 2018;12: 101. doi:10.3389/fnbeh.2018.00101

48. Sarid S, Naor H, Asfur M, Khokhlova IS, Krasnov BR, Kotler BP, et al. Free-living gerbils with higher testosterone take fewer risks. Physiol Behav. 2023;269: 114277. doi:10.1016/j.physbeh.2023.114277

49. Ronay R, van der Meij L, Oostrom JK, Pollet TV. No Evidence for a Relationship Between Hair Testosterone Concentrations and 2D:4D Ratio or Risk Taking. Front Behav Neurosci. 2018;12. doi:10.3389/fnbeh.2018.00030

50. Stanton SJ, Mullette-Gillman OA, McLaurin RE, Kuhn CM, LaBar KS, Platt ML, et al. Low- and high-testosterone individuals exhibit decreased aversion to economic risk. Psychol Sci. 2011;22: 447–453. doi:10.1177/0956797611401752

51. Kluen LM, Agorastos A, Wiedemann K, Schwabe L. Cortisol boosts risky decision-making behavior in men but not in women. Psychoneuroendocrinology. 2017;84: 181–189. doi:10.1016/j.psyneuen.2017.07.240

52. Lighthall NR, Mather M, Gorlick MA. Acute stress increases sex differences in risk seeking in the balloon analogue risk task. PloS One. 2009;4: e6002. doi:10.1371/journal.pone.0006002

53. Ceccato S, Kudielka BM, Schwieren C. Increased Risk Taking in Relation to Chronic Stress in Adults. Front Psychol. 2016;6. doi:10.3389/fpsyg.2015.02036

54. Mehta PH, Josephs RA. Testosterone and cortisol jointly regulate dominance: evidence for a dual-hormone hypothesis. Horm Behav. 2010;58: 898–906. doi:10.1016/j.yhbeh.2010.08.020

55. Laudenslager ML, Jorgensen MJ, Grzywa R, Fairbanks LA. A novelty seeking phenotype is related to chronic hypothalamic-pituitary-adrenal activity reflected by hair cortisol. Physiol Behav. 2011;104: 291–295. doi:10.1016/j.physbeh.2011.03.003

56. Laudenslager ML, Jorgensen MJ, Fairbanks LA. Developmental patterns of hair cortisol in male and female nonhuman primates: Lower hair cortisol levels in vervet males emerge at puberty. Psychoneuroendocrinology. 2012;37: 1736–1739. doi:10.1016/j.psyneuen.2012.03.015

57. Fernandez-Lazaro G, Latorre R, Alonso-Garcia E, Núñez IB. Nonhuman primate welfare: Can there be a relationship between personality, lateralization and physiological indicators? Behav Processes. 2019;166: 103897.

58. Archard GA, Braithwaite VA. The importance of wild populations in studies of animal temperament. J Zool. 2010;281: 149–160.

59. Cheney DL, Seyfarth RM. Selective Forces Affecting the Predator Alarm Calls of Vervet Monkeys. Behaviour. 1981;76: 25–61.

60. Seyfarth RM, Cheney DL, Marler P. Monkey Responses to Three Different Alarm Calls: Evidence of Predator Classification and Semantic Communication. Science. 1980;210: 801–803. doi:10.1126/science.7433999

61. Fairbanks LA, Jorgensen MJ, Huff A, Blau K, Hung Y-Y, Mann JJ. Adolescent impulsivity predicts adult dominance attainment in male vervet monkeys. Am J Primatol. 2004;64: 1–17. doi:10.1002/ajp.20057

62. Fairbanks LA, Bailey JN, Breidenthal SE, Laudenslager ML, Kaplan JR, Jorgensen MJ. Environmental stress alters genetic regulation of novelty seeking in vervet monkeys. Genes Brain Behav. 2011;10: 683–688. doi:10.1111/j.1601-183X.2011.00707.x

63. McGuire MT, Raleigh MJ, Pollack DB. Personality features in vervet monkeys: The effects of sex, age, social status, and group composition. Am J Primatol. 1994;33: 1–13. doi:10.1002/ajp.1350330102

64. Nowak K, Richards SA, Le Roux A, Hill RA. Influence of live-capture on risk perceptions of habituated samango monkeys. J Mammal. 2016;97: 1461–1468. doi:10.1093/jmammal/gyw083

65. Wasserman MD, Chapman CA, Milton K, Goldberg TL, Ziegler TE. Physiological and Behavioral Effects of Capture Darting on Red Colobus Monkeys (Procolobus rufomitratus) with a Comparison to Chimpanzee (Pan troglodytes) Predation. Int J Primatol. 2013;34: 1020–1031. doi:10.1007/s10764-013-9711-y

66. Fairbanks LA, Bailey JN, Breidenthal SE, Laudenslager ML, Kaplan JR, Jorgensen MJ. Environmental stress alters genetic regulation of novelty seeking in vervet monkeys. Genes Brain Behav. 2011;10: 683–688. doi:10.1111/j.1601-183X.2011.00707.x

67. Gesquiere LR, Learn NH, Simao MCM, Onyango PO, Alberts SC, Altmann J. Life at the Top: Rank and Stress in Wild Male Baboons. Science. 2011;333: 357–360. doi:10.1126/science.1207120

68. Laubi B, Glauser G, Willems E, Burkart J, Schaik C. Insights into Hair Steroid Dynamics in Male and Female Common Marmosets (Callithrix Jacchus) Using a Novel Hplc-Ms/Ms Assay. 2025. doi:10.2139/ssrn.5124513

69. Mehlman PT, Higley J, Faucher I, Lilly A, Taub DM, Vickers J, et al. Low CSF 5-HIAA concentrations and severe aggression and impaired impulse control in nonhuman primates. Am J Psychiatry. 1994;151: 1485–1491.

70. Laskowski KL, Chang C-C, Sheehy K, Aguiñaga J. Consistent Individual Behavioral Variation: What Do We Know and Where Are We Going? Annu Rev Ecol Evol Syst. 2022;53: 161–182. doi:10.1146/annurev-ecolsys-102220-011451

71. Mazza V, Jacob J, Dammhahn M, Zaccaroni M, Eccard JA. Individual variation in cognitive style reflects foraging and anti-predator strategies in a small mammal. Sci Rep. 2019;9: 10157. doi:10.1038/s41598-019-46582-1

72. Bell AM, Hankison SJ, Laskowski KL. The repeatability of behaviour: a meta-analysis. Anim Behav. 2009;77: 771–783. doi:10.1016/j.anbehav.2008.12.022

73. Pillay KR, Streicher JP, Downs CT. Trends in vervet monkey admissions to a wildlife rehabilitation centre: a reflection of human-wildlife conflict in an urban-forest mosaic landscape. Mamm Biol. 2024;104: 707–723. doi:10.1007/s42991-024-00447-x

74. Brett FL, Turner TR, Jolly CJ, Cauble RG. Trapping Baboons and Vervet Monkeys from Wild, Free-Ranging Populations. J Wildl Manag. 1982;46: 164. doi:10.2307/3808419

75. Etting SF, Isbell LA, Grote MN. Factors increasing snake detection and perceived threat in captive rhesus macaques (Macaca mulatta). Am J Primatol. 2014;76: 135–145.

76. Quintero F, Touitou S, Magris M, Zuberbühler K. An Audience Effect in Sooty Mangabey Alarm Calling. Front Psychol. 2022;Volume 13-2022. Available: https://www.frontiersin.org/journals/psychology/articles/10.3389/fpsyg.2022.816744

77. Schad L, van de Waal E, Fischer J. Anti-Snake Behavior and Snake Discrimination in Vervet Monkeys. Ethology. 2025;131: e13541. doi:10.1111/eth.13541

78. Etting SF, Isbell LA. Rhesus Macaques (Macaca mulatta) Use Posture to Assess Level of Threat From Snakes. Ethology. 2014;120: 1177–1184.

79. Opreni F, van de Waal E, Harrison R. Uncovering variation in social tolerance among wild vervet monkeys through a novel co-feeding paradigm. bioRxiv. 2025; 2025.08.03.668336. doi:10.1101/2025.08.03.668336

80. Godoy I, Korsten P, Perry SE. Genetic, maternal, and environmental influences on sociality in a pedigreed primate population. Heredity. 2022;129: 203–214. doi:10.1038/s41437-022-00558-6

81. Watson SK, Vale GL, Hopper LM, Dean LG, Kendal RL, Price EE, et al. Chimpanzees demonstrate individual differences in social information use. Anim Cogn. 2018;21: 639–650. doi:10.1007/s10071-018-1198-7

82. Cheney DL, Seyfarth RM. How Monkeys See the World: Inside the Mind of Another Species. University of Chicago Press; 1990. Available: https://books.google.ch/books?id=QbWTd85eHUgC

83. Laviola G, Macrì S, Morley-Fletcher S, Adriani W. Risk-taking behavior in adolescent mice: psychobiological determinants and early epigenetic influence. Neurosci Biobehav Rev. 2003;27: 19–31. doi:10.1016/s0149-7634(03)00006-x

84. Rosati AG, Emery Thompson M, Atencia R, Buckholtz JW. Distinct developmental trajectories for risky and impulsive decision-making in chimpanzees. J Exp Psychol Gen. 2023;152: 1551–1564. doi:10.1037/xge0001347

85. Josef AK, Richter D, Samanez-Larkin GR, Wagner GG, Hertwig R, Mata R. Stability and change in risk-taking propensity across the adult life span. J Pers Soc Psychol. 2016;111: 430–450. doi:10.1037/pspp0000090

86. Willoughby T, Heffer T, Good M, Magnacca C. Is adolescence a time of heightened risk taking? An overview of types of risk-taking behaviors across age groups. Dev Rev. 2021;61: 100980. doi:10.1016/j.dr.2021.100980

87. Fairbanks LA, McGuire MT. Maternal protectiveness and response to the unfamiliar in vervet monkeys. Am J Primatol. 1993;30: 119–129. doi:10.1002/ajp.1350300204

88. Fairbanks LA, Jorgensen MJ. Objective Behavioral Tests of Temperament in Nonhuman Primates. In: Weiss A, King JE, Murray L, editors. Personality and Temperament in Nonhuman Primates. New York, NY: Springer; 2011. pp. 103–127. doi:10.1007/978-1-4614-0176-6_5

89. Ellington L, Mercier S, Motes-Rodrigo A, van de Waal E, Forss S. Urbanization does not increase “object curiosity” in vervet monkeys, but semi-urban individuals selectively explore food-related anthropogenic items. Curr Zool. 2024;70: 383–393. doi:10.1093/cz/zoae022

90. Laube C, van den Bos W. Hormones and Affect in Adolescent Decision Making. Recent Developments in Neuroscience Research on Human Motivation. Emerald Group Publishing Limited; 2016. p. 0. doi:10.1108/S0749-742320160000019013

91. Jolles JW, Boogert NJ, van den Bos R. Sex differences in risk-taking and associative learning in rats. R Soc Open Sci. 2015;2: 150485. doi:10.1098/rsos.150485

92. Ren S, Liu S, Sun W, Gao L, Ren L, Liu J, et al. Consistent Individual Differences Drive Collective Movements in a Tibetan Macaque Group. Animals. 2024;14: 1476. doi:10.3390/ani14101476

93. Neumann C, Agil M, Widdig A, Engelhardt A. Personality of Wild Male Crested Macaques (Macaca nigra). PLoS ONE. 2013;8: e69383. doi:10.1371/journal.pone.0069383

94. Baker MD, Maner JK. Risk-taking as a situationally sensitive male mating strategy. Evol Hum Behav. 2008;29: 391–395. doi:10.1016/j.evolhumbehav.2008.06.001

95. Tankink JA, van de Waal E, Bshary R, van Schaik CP. Leadership under risk: male vervet monkey’s roles in group progression across high-risk terrain. bioRxiv. 2025; 2025.11.21.689472. doi:10.1101/2025.11.21.689472

96. Bell AM, Hankison SJ, Laskowski KL. The repeatability of behaviour: a meta-analysis. Anim Behav. 2009;77: 771–783. doi:10.1016/j.anbehav.2008.12.022

97. Cheney DL, Seyfarth RM. Vervet Monkey Alarm Calls: Manipulation through Shared Information? Behaviour. 1985;94: 150–166.

98. van Schaik CP, Bshary R, Wagner G, Cunha F. Male anti-predation services in primates as costly signalling? A comparative analysis and review. Ethology. 2022;128: 1–14. doi:10.1111/eth.13233

99. Baldellou M, Peter Henzi S. Vigilance, predator detection and the presence of supernumerary males in vervet monkey troops. Anim Behav. 1992;43: 451–461. doi:10.1016/S0003-3472(05)80104-6

100. Majolo B, Lehmann J, de Bortoli Vizioli A, Schino G. Fitness-related benefits of dominance in primates. Am J Phys Anthropol. 2012;147: 652–660. doi:10.1002/ajpa.22031

101. Abbott DH, Keverne EB, Bercovitch FB, Shively CA, Mendoza SP, Saltzman W, et al. Are subordinates always stressed? a comparative analysis of rank differences in cortisol levels among primates. Horm Behav. 2003;43: 67–82. doi:10.1016/S0018-506X(02)00037-5

102. Amici F, Widdig A, MacIntosh AJJ, Francés VB, Castellano-Navarro A, Caicoya AL, et al. Dominance style only partially predicts differences in neophobia and social tolerance over food in four macaque species. Sci Rep. 2020;10: 22069. doi:10.1038/s41598-020-79246-6

103. Huang P, Arlet ME, Balasubramaniam KN, Beisner BA, Bliss-Moreau E, Brent LJN, et al. Relationship between dominance hierarchy steepness and rank-relatedness of benefits in primates. Behav Ecol. 2024;35: arae066. doi:10.1093/beheco/arae066

104. Schad L, Dongre P, van de Waal E, Fischer J. Loud Call Production in Male Vervet Monkeys (Chlorocebus pygerythrus) Varies with Season and Signaller Rank. Int J Primatol. 2025;46: 538–555. doi:10.1007/s10764-024-00475-x

105. Garber PA, McKenney A, Bartling-John E, Bicca-Marques JC, Fuente MFD la, Abreu F, et al. Life in a harsh environment: the effects of age, sex, reproductive condition, and season on hair cortisol concentration in a wild non-human primate. PeerJ. 2020;8: e9365. doi:10.7717/peerj.9365

106. Raulo A, Dantzer B. Associations between glucocorticoids and sociality across a continuum of vertebrate social behavior. Ecol Evol. 2018;8: 7697–7716. doi:10.1002/ece3.4059

107. Russell E, Koren G, Rieder M, Van Uum S. Hair cortisol as a biological marker of chronic stress: Current status, future directions and unanswered questions. Psychoneuroendocrinology. 2012;37: 589–601. doi:10.1016/j.psyneuen.2011.09.009

108. Yamanashi Y. Is Hair Cortisol Useful for Animal Welfare Assessment? Review of Studies in Captive Chimpanzees. Aquat Mamm. 2018;44: 201–210. doi:10.1578/am.44.2.2018.201

109. Cunningham EP, Unwin S, Setchell JM. Darting Primates in the Field: A Review of Reporting Trends and a Survey of Practices and Their Effect on the Primates Involved. Int J Primatol. 2015;36: 911–932. doi:10.1007/s10764-015-9862-0

110. Koren L, Bryan H, Matas D, Tinman S, Fahlman Å, Whiteside D, et al. Towards the validation of endogenous steroid testing in wildlife hair. J Appl Ecol. 2019;56: 547–561.

111. Koren L, Geffen E. Androgens and social status in female rock hyraxes. Anim Behav. 2009;77: 233–238. doi:10.1016/j.anbehav.2008.09.031

112. Friard O, Gamba M. BORIS: a free, versatile open-source event-logging software for video/audio coding and live observations. Methods Ecol Evol. 2016;7: 1325–1330. doi:10.1111/2041-210X.12584

113. Altmann J. Observational study of behavior: sampling methods. Behaviour. 1974;49: 227–267. doi:10.1163/156853974×00534

114. Neumann C, Duboscq J, Dubuc C, Ginting A, Maulana A, Agil M, et al. Assessing dominance hierarchies: Validation and advantages of progressive evaluation with Elo-rating. Anim Behav. 2011;82: 911–921. doi:10.1016/j.anbehav.2011.07.016

115. R Core Team. R: A language and environment for statistical computing. R Foundation for Statistical Computing, Vienna, Austria; 2021. Available: https://www.R-project.org/

116. RStudio Team. RStudio: Integrated Development for R. RStudio. PBC.; 2022. Available: http://www.rstudio.com/

117. Revelle W. psych: Procedures for Psychological, Psychometric, and Personality Research. Evanston, Illinois: Northwestern University; 2025. Available: https://CRAN.R-project.org/package=psych

118. Morton FB, Altschul D. Data reduction analyses of animal behaviour: avoiding Kaiser’s criterion and adopting more robust automated methods. Anim Behav. 2019;149: 89–95. doi:10.1016/j.anbehav.2019.01.003

119. Stoffel M, Schielzeth H. rptR: Repeatability estimation and variance decomposition by generalized linear mixed-effects models. Methods Ecol Evol. 2017;8. doi:10.1111/2041-210X.12797

120. Bates D, Mächler M, Bolker B, Walker S. Fitting Linear Mixed-Effects Models Using lme4. J Stat Softw. 2015;67: 1–48. doi:10.18637/jss.v067.i01

121. Brooks ME, Kristensen K, Benthem KJ van, Magnusson A, Berg CW, Nielsen A, et al. glmmTMB Balances Speed and Flexibility Among Packages for Zero-inflated Generalized Linear Mixed Modeling. R J. 2017;9: 378–400. doi:10.32614/RJ-2017-066

122. Fox J, Weisberg S. An R Companion to Applied Regression. Third. Thousand Oaks CA: Sage; 2019. Available: https://www.john-fox.ca/Companion/

123. Hartig F. DHARMa: Residual Diagnostics for Hierarchical (Multi-Level / Mixed) Regression Models. 2024. Available: http://florianhartig.github.io/DHARMa/

124. Lüdecke D, Ben-Shachar MS, Patil I, Waggoner P, Makowski D. performance: An R Package for Assessment, Comparison and Testing of Statistical Models. J Open Source Softw. 2021;6: 3139. doi:10.21105/joss.03139

