## Supplemental files for "Who takes the risk to forage? Testing consistent inter-individual behavioural variation in wild vervet monkeys (*Chlorocebus pygerythrus*)"

Short title: Individual variation in risk-taking behaviour in wild vervet monkeys

des Pallières Claire Gauquelin, Belli Elena, Aguilera Fanny, Halbwax Michel, Šlipogor Vedrana, van de Waal Erica, Koren Lee, Matas Devorah, Canteloup Charlotte

<table>
<tr><th colspan="4">Effects of socio-demographic factors on risky behaviours</th></tr>
<tr><td>Approach model</td><td>Alarm call</td><td>Scans within 1m to 5m of the corn</td><td>Scans within 1m of the corn</td></tr>
<tr><td>GLMM_2:<br>GLMM Poisson (link 'log')</td><td>ZIP_1:<br>Zero-inflated GLMM Poisson (link 'log')</td><td>ZINB_1:<br>Zero-inflated GLMM binomial (link 'logit')</td><td>GLMM_1: GLMM binomial (link 'logit')</td></tr>
<tr><td>Counts with an offset for trial duration</td><td>Counts with an offset for trial duration</td><td>Proportion of successes versus failures</td><td>Proportion of successes versus failures</td></tr>
<tr><td>Age +<br>Sex +<br>Rank + Experiment type</td><td>Age +<br>Sex +<br>Rank + Experiment type</td><td>Age +<br>Sex +<br>Rank + Experiment type</td><td>Age +<br>Sex +<br>Rank + Experiment type</td></tr>
<tr><td>(1 | Group/ID) + (1 | Experiment number)</td><td>(1 | Group/ID) + (1 | Experiment number)</td><td>(1 | Group/ID) + (1 | Experiment number)</td><td>(1 | Group/ID) + (1 | Experiment number)</td></tr>
<tr><td>7 | 222 | 1910</td><td>7 | 222 | 1910</td><td>7 | 222 | 1910</td><td>7 | 222 | 1910</td></tr>
<tr><td>(i) Juveniles approach the model more often than adults<br>(ii) Males approach the model more often than females<br>(iii) No effect of rank<br>(iv) More model approaches should occur in snake > darting > eagle</td><td>(i) No strong effect of age<br>(ii) No strong effect of sex<br>(iii) No strong effect of rank<br>(iv) Most alarm calls occur in snake > eagle > darting</td><td>(i) No strong effect of age<br>(ii) No strong effect of sex<br>(iii) No strong effect of rank<br>(iv) Individuals are more likely to be within 5m of corn during darting > snake > eagle</td><td>(i) Juveniles spend more time in 1m of corn than adults<br>(ii) Males spend more time in 1m of corn than females<br>(iii) No strong effect of rank<br>(iv) Individuals are more likely to be in 1m of corn during darting > snake > eagle</td></tr>
<tr><td>(i) As predicted<br>(ii) No effect of sex<br>(iii) As predicted<br>(iv) More approaches in snake > eagle = darting</td><td>(i) As predicted<br>(ii) Males alarm call at higher rate than females<br>(iii) High-rank individuals are more likely to alarm call<br>(iv) As predicted</td><td>(i) Juveniles spend more time within 5m<br>(ii) As predicted<br>(iii) High-ranking individuals are more likely to be within 5m<br>(iv) No effect of Experiment</td><td>(i) As predicted<br>(ii) As predicted<br>(iii) High-rank individuals are more likely to be within 1m<br>(iv) Individuals are more likely to be within 1m in snake compared to darting, with no significant difference between eagle-snake or eagle-darting</td></tr>
</table>

| Question | Effects of hormonal levels on risky behaviours | | Effects of socio-demographic factors on hormones | |
|---|---|---|---|---|
| **Dependent Variable** | Alarm call | Scans within 1m of the corn | Cortisol | Testosterone |
| **Model** | GLMM_4: GLMM Poisson (link 'log') | GLMM_3: GLMM binomial (link 'logit') | LM_2: LM | LM_1: LM |
| **Data Type** | Counts with an offset for trial duration | Proportion of successes versus failures | Continuous (pg/mg hair) | Continuous (pg/mg hair) |
| **Fixed predictors** | Testosterone + Cortisol + Experiment | Testosterone + Cortisol + Experiment | Sex + Rank | Sex + Rank |
| **Random effects** | (1 \| ID) | (1 \| ID) | None | None |
| **Nb. Groups \| Nb. Individuals \| Nb.** | 7 \| 35 \| 297 | 7 \| 35 \| 297 | 7 \| 37 \| 37 | 7 \| 37 \| 37 |
| **Hypotheses** | (i) Individuals with high levels of testosterone alarm call more<br>(ii) Individuals with low levels of cortisol alarm call more<br>(iii) Individuals are more likely to alarm call during snake > eagle > darting | (i) Individuals with high levels of testosterone are more risk prone<br>(ii) Individuals with low levels of cortisol are more risk prone<br>(iii) Individuals are more likely to be in 1m of corn during darting > snake > eagle | (i) Males have lower cortisol levels than females<br>(ii) High ranking individuals have lower cortisol levels | (i) Males have higher testosterone levels than females<br>(ii) No strong effect of rank |
| **Results** | (i) No effect<br>(ii) No effect<br>(iii) Individuals are more likely to alarm call during snake > eagle > darting | (i) No effect<br>(ii) No effect<br>(iii) Individuals are least likely to be within 1m in darting | (i) As predicted<br>(ii) No effect | (i) Non-significant trend in the predicted direction<br>(ii) As predicted |

**S1 Table.** Summary of the models used for our analysis. Response variables included scans within 1 m and 5 m of the corn (binomial models), alarm calls and approach behaviour (count models with log link), with trial duration included as an offset where appropriate. Fixed effects were age, sex, rank and experiment type; random effects were group and individual identity nested within group (Group/ID), with experiment number included where relevant. Sample sizes are given for each model. Hypotheses and corresponding results are summarised.

**S1 Text.** Description of models used to investigate compare the control condition to our main experimental conditions.

We ran three models to test for the effect of experiment type (including the turtle control) on specific behaviours of interest (number of scans spent within 1m of the corn, rate of alarm calling, and rate of vigilance). As we only carried out one trial per group for the control condition, we made a subset of data that included only the first trial of each experiment type (turtle, snake, eagle, darting). We ran a GLMM with a binomial distribution and logit link function with "scans within 1m of the corn" as dependent variable. This was modelled as the number of 'successes' (i.e., number of scans where an ID was observed within 1m of the corn) versus 'failures' (i.e., number of scans where that ID was not within 1m of the corn), allowing for a proportional outcome. Due to zero-inflated data and poor residuals plots, we fitted negative binomial models for the dependent variables "alarm calling" (ZIP_3) and "vigilance" (ZIP_4). These variables were modelled as the occurrence (count) of the behaviour (i.e., number of alarm calls given) and included an offset for the trial duration to account for different observation lengths. In all three models, experiment type was added as a fixed effect, and a random effect of ID nested within Group. In BIN_2, ID nested within group was added as a random effect.

| Groups | October | November | December | January | February | March |
|---|---|---|---|---|---|---|
| AK | **20** (AM: 3, AF: 7, JM:6, JF: 4 , UNJ: 0) | **21** (AM: 3; AF: 7; JM: 7; JF: 4; UNKJ: 0) | **21** (AM: 3; AF: 7; JM: 7; JF: 4; UNKJ: 0) | **21** (AM: 3; AF: 7; JM: 7; JF: 4; UNKJ: 0) | **20** (AM: 3; AF: 7; JM: 7; JF: 3; UNKJ: 0) | **19** (AM: 2; AF: 7; JM: 7; JF: 3; UNKJ: 0) |
| BD | **72** (21, 23, 15, 13, 0) | **70** (19; 23; 15; 13; 0) | **69** (19; 23; 14; 13; 0) | **66** (16; 23; 14; 13; 0) | **43** (8; 12; 13; 10; 0) | **44** (9; 12; 13; 10; 0) |
| CR | **28** (7, 9, 6, 6, 0) | **27** (6; 9; 6; 6; 0) | **27** (6; 9; 6; 6; 0) | **26** (5; 9; 6; 6; 0) | **25** (4; 9; 6; 6; 0) | **25** (4; 9; 6; 6; 0) |
| IF | **23** (5; 7; 4; 4; 3) | **NA** | **NA** | **NA** | **NA** | **NA** |
| KB | **22** (4; 6; 4; 8; 0) | **19** (3; 6; 4; 6; 0) | **19** (3; 6; 4; 6; 0) | **19** (3; 6; 4; 6; 0) | **19** (3; 6; 4; 6; 0) | **19** (3; 6; 4; 6; 0) |
| LT | **34** (7; 12; 4; 11; 0) | **35** (7; 12; 4; 12; 0) | **36** (7; 12; 4; 12; 1) | **36** (6; 12; 3; 11; 3) | **36** (6; 12; 3; 11; 3) | **36** (6; 12; 3; 11; 3) |
| NH | **45** (9; 16; 9; 11; 0) | **41** (6; 15; 9; 11; 0) | **41** (6; 15; 9; 11; 0) | **33** (6; 9; 8; 10; 0) | **32** (7; 8; 7; 10; 0) | **32** (7; 8; 7; 10; 0) |
| Total | **242** (55; 80; 48; 55; 4) | **213** (44; 72; 45; 52; 0) | **213** (44; 72; 44; 52; 1) | **200** (39; 66; 42; 50; 3) | **176** (31; 54; 42; 46; 3) | **176** (31; 54; 42; 46; 3) |

**S2 Table.** Demographic composition of the seven habituated study groups from October 2023 to April 2024. Group composition is expressed as the total number of individuals (in bold), and the detail per age and sex categories is provided within brackets (n adult male (AM); n adult female (AF); n juvenile male (JM); n juvenile female (JF); n unsexed juveniles (UNJ)).

**S2 Text.** Results from our analysis of the control turtle condition.
Experiment type had a significant effect on "scans within 1m of the corn" and significantly improved model fit when compared to a null model ($\chi 2 = 93.079$; df = 3; $p < 0.0001$; N = 444). When compared to the turtle condition, individuals spent significantly fewer scans within 1m of the corn in the eagle experiments (Estimate = -1.65; SE = 0.22; z = -7.56; $p < 0.0001$), snake experiments (Estimate = -1.45; SE = 0.19; z = -7.75; $p < 0.0001$), and darting experiments (Estimate = -1.85; SE = 0.19; z = -9.66; $p < 0.0001$). The random effect of ID nested within group had a high variance of $\sigma^2 = 2.16$ (SD = 1.47) and group alone had a variance of $\sigma^2 = 0.91$ (SD = 0.95). The marginal $R^2$ was 0.096, indicating that experiment type alone explained approximately 9.6% of the variance in the amount of time spent within 1m of the corn. The conditional $R^2$ was 0.915, suggesting that the full model—including the random effect of ID nested within group—explained approximately 91.5% of the variance. Experiment type had a significant effect on alarm calling and its inclusion significantly improved model fit ($\chi 2 = 62.672$; df = 3; $p < 0.0001$; N = 444). When compared to the turtle condition, there was a significantly higher rate of alarm calling in the snake experiments (Estimate = 4.25; SE = 1.27; z = 3.36; p = 0.001), but there was no significant differences in the eagle experiments (Estimate = 1.21; SE = 1.31; z = 0.93, $p > 0.05$), or in the darting (Estimate = -2.17; SE = 1.51; z = -1.44; p = 0.149). The random effect of ID nested within group had a high variance of $\sigma^2 = 2.11$ (SD = 1.45) and group alone had a variance of $\sigma^2 = 0.23$ (SD = 0.48). The marginal $R^2$ was 0.392, indicating that experiment alone explained approximately 39.2% of the variance in the amount of alarm calling. The conditional $R^2$ was 0.545, suggesting that the full model—including the random effect of ID nested within group—explained approximately 54.5% of the variance. Experiment had a significant effect on number of vigilance behaviours ($\chi 2 = 70.669$; df = 3; $p < 0.0001$; N = 444). When compared to the turtle condition, there was significantly lower rates of vigilance in eagle experiments (Estimate = -1.48; SE = 0.37; z = -4.02; $p < 0.0001$), and in darting experiments (Estimate= -2.88, SE = 0.36; z = -7.99; $p < 0.0001$), but there was no significant difference in the snake experiments (Estimate = -0.34; SE = 0.32; z = -1.07; p = 0.283). The random effect of ID nested within group had a low variance of $\sigma^2 = 0.41$ (SD= 0.64) and group alone had a variance of $\sigma^2 = 0.19$ (SD = 0.44). The marginal $R^2$ was 0.143, indicating that experiment alone explained approximately 14.3% of the variance in the amount of vigilance. The conditional $R^2$ was 0.218, suggesting that the full model including the random effect of ID nested within group—explained approximately 21.8% of the variance.

| Behaviour | Experiment | Type | Category | Description |
| --- | --- | --- | --- | --- |
| **Scans within 1m of the corn** | All | Event | Distance | For the scan, individual is within arms' reach of the corn. |
| **Scans within 1 to 5m of the corn** | All | Event | Distance | For the scan, individual is within 5m of the corn. |
| **Scans within 5m to 10m of the corn** | All | Event | Distance | For the scan, individual is within 6 to 10m of corn. |
| **Scans in >10m from the corn** | All | Event | Distance | For the scan, individual is visible outside of 10m from the corn. |
| **Approach the corn** | All | Event | Distance | Number of times an individual arrives within 1m of the corn during the experiment. |
| **Absent** | All | Event | Distance | Individual was observed in the group that day, but not visible during the experiment. |
| **Time spent at the corn** | All | State | Feeding behaviour | Individual is within arm's reach of the corn, this includes actively feeding or not. |
| **Allo-groom** | All | Event | Affiliative | Individual combs through the hair of a conspecific to remove dead skin and parasites. |
| **Vocalise** | All | Event | Affiliative | Individual produces any call that is not an alarm call or a scream. |
| **Stare attack** | All | Event | Agonistic | Individual raises its eyebrows, opening its eyes wide and moves is body towards an individual. |
| **Physical attack** | All | Event | Agonistic | Individual physically interacts with another in an agonistic manner (e.g. hit, bite, fight). |
| **Head-bob** | All | Event | Agonistic | Individual moves its head up and down repeatedly at another individual. |
| **Jump aside** | All | Event | Agonistic | Individual jumps in the air to avoid another one. |
| **Scream** | All | Event | Agonistic | Individual produces a distress call usually during agonistic situations. |
| **Chase** | All | Event | Agonistic | Individual runs after another fleeing individual. |
| **Flee from individual** | All | Event | Agonistic | Individual runs away from an aggressor. |
| **Alarm call** | All | Event | Vigilance | Individual produces an alarm call. |
| **Vigilant** | All | Event | Vigilance | Individual stands bipedally to scan its surroundings and/or the model. |
| **Flee from the model** | All | Event | Vigilance | Individual runs rapidly away from the experimental area. |
| **Head-bob the model** | All | Event | Vigilance | Individual moves its head up and down repeatedly at the model. |
| **Approach the model** | All | Event | Vigilance | Individual directly approaches and inspects the model from within 5m. |
| **Inspect the hide** | Darting | Event | Vigilance | Individual examines carefully the hide for at least 3s. |
| **Approach the hide** | Darting | Event | Vigilance | Individual moves toward the hide. |
| **Auto-groom** | All | Event | Self-directed | Individual combs through its own hair, removing dead skin and/or parasites. |

| | | | | |
|---|---|---|---|---|
| **Self-scratch** | All | Event | Self-directed | Individual scratches its body repeatedly using hands or feet. |
| **Yawn** | All | Event | Self-directed | Individual opens its mouth wide displaying its teeth. |

**S3 Table**. Ethogram of the behavioural variables coded during the video analysis of our experiments. The 'Experiment' column indicates whether the variable was used in all experiments, only in the 'Darting' experiment, or only in the 'Predator' experiments. The 'Type' and 'Category' columns respectively display the behaviour as an event or point variable and one of the eight categories to which it belongs.

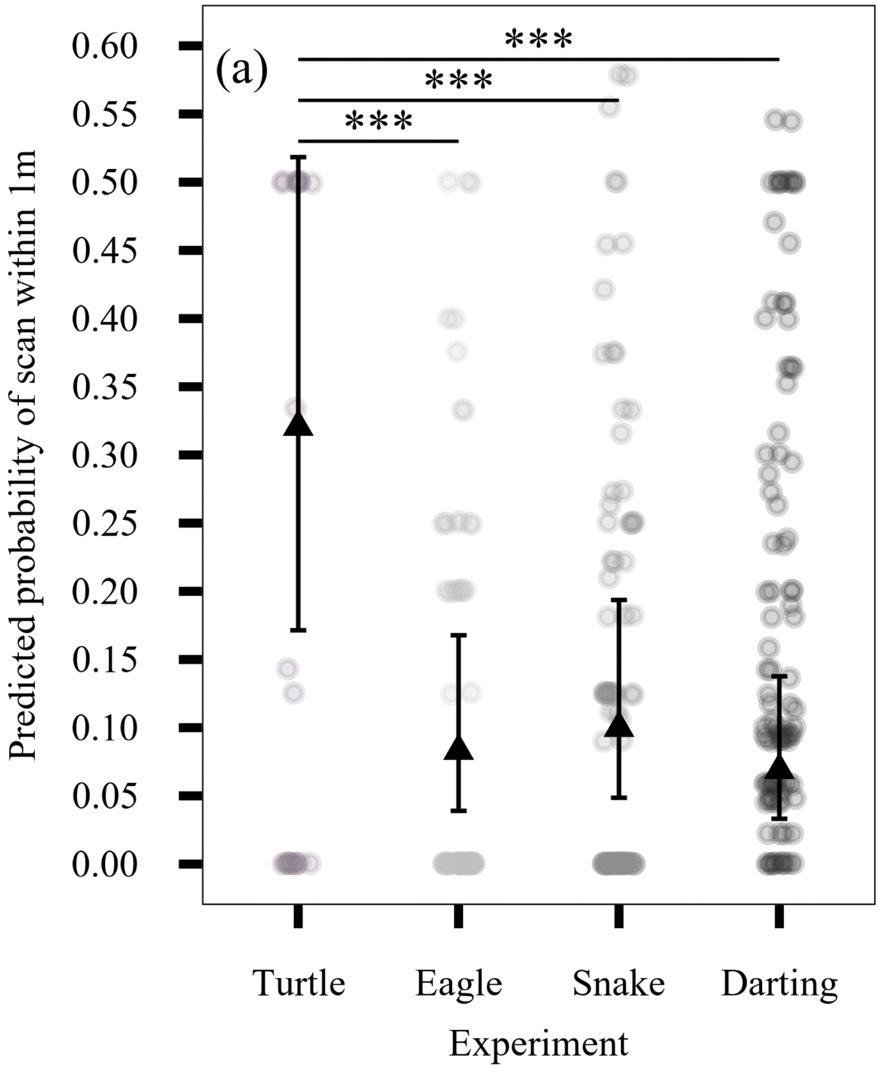


**S1a Fig**. **Predicted probability of individuals being within 1 m of the corn during a scan across experiment types, including the turtle control**.
Grey points represent the probability for one individual being within 1 m of the corn during a scan. The central triangles represent model-estimated means and error bars indicate 95% confidence intervals. Significance levels are indicated as follows:
* $p < 0.05$; ** $p < 0.01$; *** $p < 0.005$.

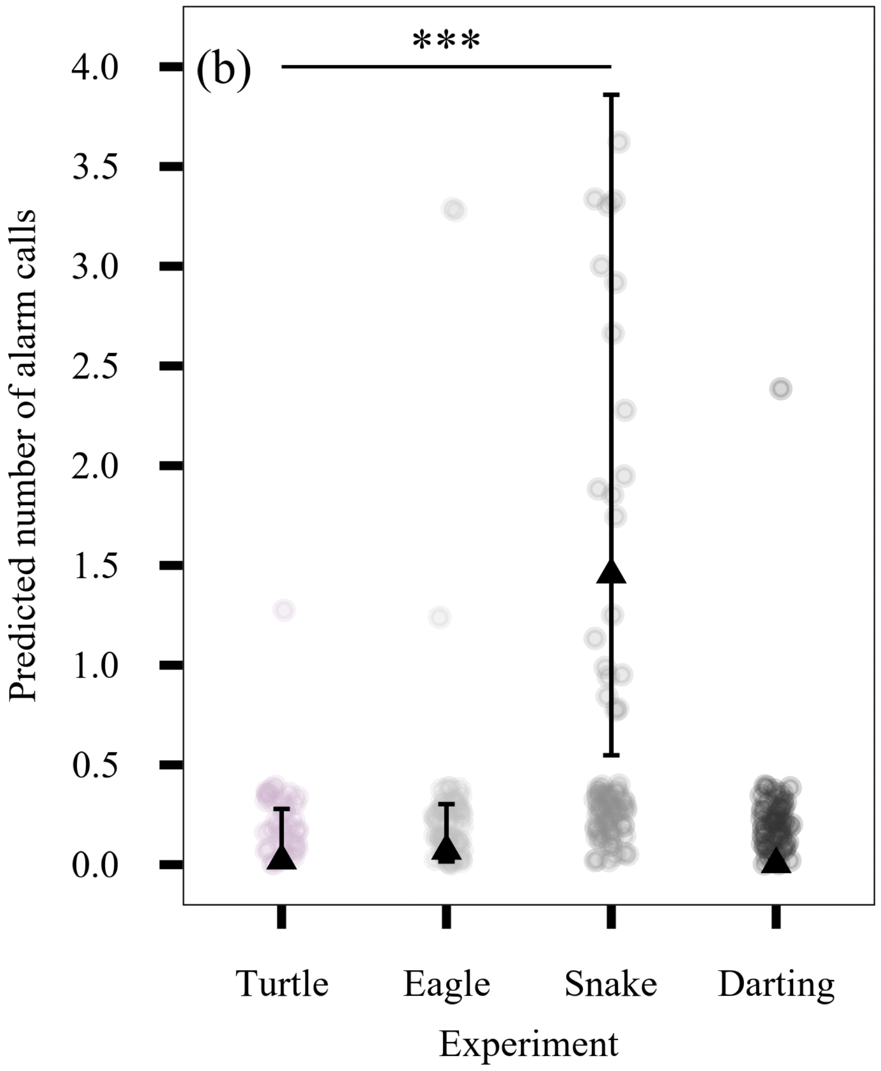


**S1b Fig. Predicted mean number of alarm calls given across experiment types, including the turtle control**.
Grey points represent the mean number of alarm calls given per individual. The solid lines and central triangles represent model-estimated mean rates of alarm calling among callers, with error bars denoting 95% confidence intervals. Significance levels are indicated as follows: * $p < 0.05$; ** $p < 0.01$; *** $p < 0.005$.

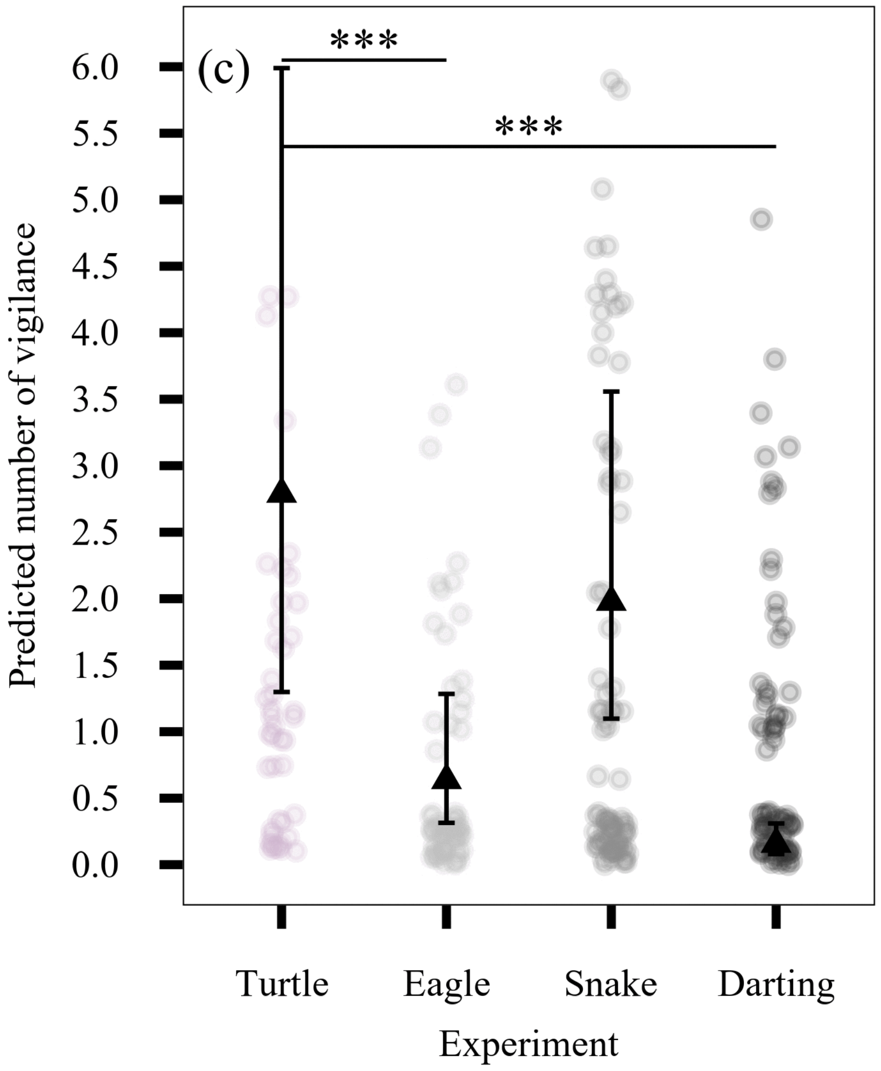


**S1c Fig. Predicted mean number of vigilance events across experiment type, including the turtle control experiment.**

Grey points represent the mean of vigilance per individual. The solid lines and central triangles represent model-estimated mean vigilance events, with error bars denoting 95% confidence intervals. Significance levels are indicated as follows: * $p < 0.05$; ** $p < 0.01$; *** $p < 0.005$.
